# Endogenous CGRP activates NRF2 signaling via non-electrophilic mechanisms

**DOI:** 10.1101/2025.04.25.650677

**Authors:** Keren Powell, Steven Wadolowski, Willians Tambo, Eric H. Chang, Daniel Kim, Asha Jacob, Daniel Sciubba, Yousef AlAbed, Ping Wang, Chunyan Li

## Abstract

The transcription factor nuclear factor erythroid 2-related factor 2 (NRF2) is crucial for regulating cellular responses to oxidative stress, making it a significant target for therapeutic interventions. While exogenous NRF2 activators offer significant therapeutic potential, their predominantly electrophilic nature poses considerable challenges for clinical use; the heightened electrophilic reactivity required to achieve therapeutic efficacy raises potential safety concerns. Calcitonin gene-related peptide (CGRP) has shown protective effects against oxidative stress and is involved in NRF2 activation; however, the underlying mechanisms are not fully understood. This study explores the mechanisms underlying endogenous CGRP-mediated NRF2 upregulation by inducing acute or chronic CGRP release through diving reflex (DR) in male Sprague-Dawley rats. Brain tissue proteomics confirmed the upregulation of NRF2-dependent antioxidant transcripts— predominantly glutathione-related genes—without concurrent elevation of oxidative stress markers in both acute and chronic CGRP exposure paradigms. CGRP potently activated NRF2 in brain and peripheral tissues, evidenced by elevated nuclear and phosphorylated NRF2, increased nuclear:cytosolic NRF2 ratios, and enhanced antioxidant gene transcription—effects substantially attenuated by CGRP antagonism. Reduced glutathione levels increased without concurrent elevations in lipid peroxidation, protein oxidation, or evidence of tissue damage, suggesting CGRP avoids side effects characteristic of electrophilic NRF2 activators. Furthermore, our findings suggest that CGRP-mediated NRF2 activation primarily occurs via non-electrophilic mechanisms, with the p62-KEAP1-NRF2 pathway predominantly active in peripheral organs (lung and kidney), and the AMPK-NRF2 pathway more pronounced in the brain, highlighting the organ-specific nature of the response. Time-dependent variations in CGRP-mediated NRF2 activation were also observed, influencing both the response to CGRP and its impact on oxidative stress resistance. These results suggest that targeting NRF2 with endogenous CGRP may offer a promising therapeutic approach for managing oxidative stress-related diseases, both acute and chronic, across multiple organs, by avoiding electrophilic stress.

**Highlights:**

- Endogenous CGRP triggers a potent and non-electrophilic activation of NRF2 signaling.
- CGRP increases reduced glutathione levels following both acute and chronic exposures, in contrast to the effects of exogenous electrophilic NRF2 activators.
- In peripheral organs, CGRP predominantly activates the KEAP1-dependent p62-KEAP1-NRF2 pathway.
- In the brain, CGRP primarily activates the KEAP1-independent AMPK-NRF2 pathway.
- CGRP exhibits time-dependent patterns, where acute exposure leads to a more significant upregulation of NRF2-targeted antioxidative gene expression and chronic exposure confers increased resistance to oxidative stress.

**Graphical Abstract:** 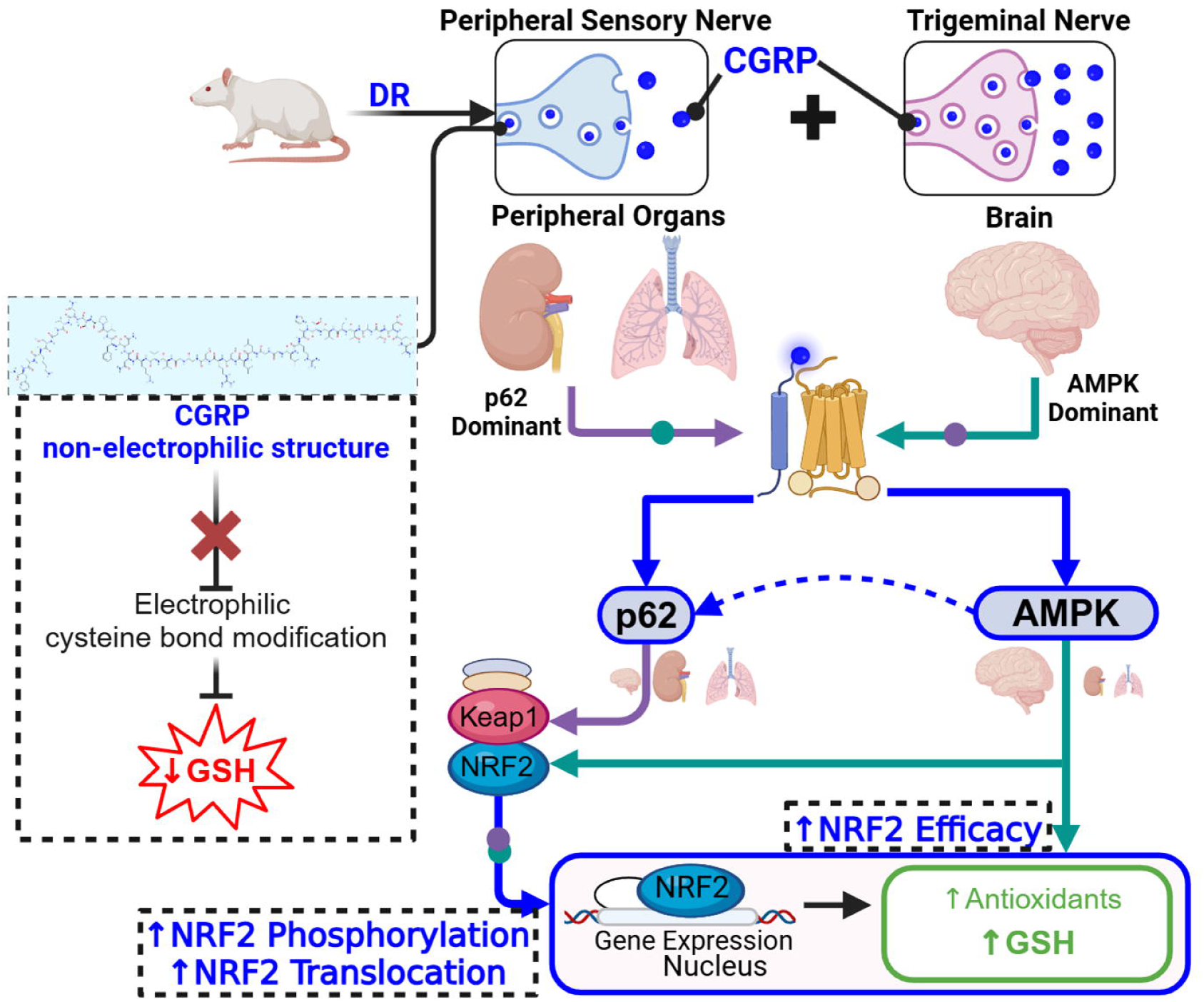

## 1. Introduction

Nuclear factor erythroid 2-related factor 2 (NRF2) is a key regulator of cellular homeostasis, particularly through its role in modulating a broad spectrum of antioxidant enzymes that detoxify and mitigate oxidative stress [1–4]. Given its central role, NRF2 is regarded as a promising therapeutic target for both chronic and acute, traumatic conditions [2,5,6]. However, while exogenous NRF2 activators hold significant therapeutic potential, the predominantly electrophilic nature of most pharmacological NRF2 activators presents several challenges to their clinical application, which is exacerbated by the increased reactivity required to achieve therapeutic efficacy. First, the depletion of intracellular glutathione reserves due to electrophilic reactions—an essential mechanism by which these activators function—can paradoxically heighten susceptibility to oxidative stress [2,7–11]. Second, chronic use may disrupt mitochondrial bioenergetics, potentially resulting in organ toxicity, particularly in conditions characterized by heightened physiological stress [5,12–15]. This underscores the need for alternative NRF2 activation strategies. Endogenous mechanisms, leveraging the body’s intrinsic processes, offer a promising approach to overcoming these limitations [12,16]. Interventions such as physical exercise, vagus nerve stimulation (VNS), electroacupuncture, and photobiomodulation have been shown to activate NRF2, without a concurrent decrease in GSH or increase in organ damage, in contrast to exogenous NRF2 activators [17–24]. However, despite the potential of these interventions, the underlying mechanisms remain unclear. Thus, exploring endogenous NRF2 activation represents a promising and underexplored research area.

Calcitonin gene-related peptide (CGRP), a neuropeptide present in sensory fibers throughout the body [25–28], is released as an endogenous protective response against oxidative stress [29–32]. CGRP plays a crucial role in maintaining antioxidative homeostasis by reducing reactive oxygen species (ROS) production in the cardiovascular and cerebrovascular systems [31,33,34] and enhancing the production of antioxidant molecules, such as endothelial nitric oxide [35] and l-citrulline [36–38]. Exogenous CGRP administration has been reported to upregulate NRF2 expression in cultured neurons [39], whereas inhibition of endogenous CGRP has been shown to suppress cerebral NRF2 upregulation induced by physical exercise [40] and reduce levels of phosphorylated NRF2 and reduced glutathione (GSH) following traumatic brain injury [41]. Although evidence suggests CGRP’s involvement in oxidative stress and NRF2 regulation, the underlying mechanisms remain unclear. This study aims to (1) investigate whether endogenous CGRP activates NRF2 signaling through mechanisms similar to, or distinct from, those of exogenous electrophilic NRF2 activators, (2) determine whether CGRP-mediated NRF2 activation elicits differential responses in the brain compared to peripheral organs, and (3) evaluate the distinct effects of acute versus chronic CGRP exposure on NRF2 signaling.

To investigate the mechanisms by which endogenous CGRP modulates NRF2, we employed a rodent model of the diving reflex (DR) [42–44], a robust physiological response that triggers systemic antioxidative responses and is linked to CGRP upregulation [45–54] (**Supplementary** Fig. 1). Acute CGRP exposure was induced through a single session of intermittent DR, while chronic exposure was achieved through daily intermittent DR sessions over four weeks. Additional controls consisted of acute and chronic DR groups treated with a CGRP antagonist to confirm CGRP as a key mediator. Brain tissue proteomics was first used to evaluate non-electrophilic NRF2 activation under acute and chronic CGRP exposure paradigms, confirming the upregulation of NRF2-dependent antioxidative transcripts—predominantly glutathione-related genes—without concomitant increases in oxidative stress markers. Following this confirmation, the NRF2 pathway analysis was extended to brain and peripheral tissues to delineate the systemic impact of endogenous CGRP signaling and assess its therapeutic potential. Total, nuclear, cytosolic, and phosphorylated NRF2 expression was quantified in the brain and peripheral organs (lung and kidney), alongside assessments of non-electrophilic NRF2 activators and electrophilic activation markers, including GSH, malondialdehyde and 4-hydroxy-2-nonenal (4-HNE) as indicators of lipid peroxidation, nitrotyrosine as a marker of protein oxidation, and peripheral organ damage. Our findings demonstrate that endogenous CGRP upregulation robustly activates NRF2 signaling via non-electrophilic pathways in an organ-specific and exposure time-dependent manner. These results suggest that modulating endogenous CGRP may serve as a potential therapeutic strategy for enhancing NRF2 activation and reducing oxidative stress while minimizing electrophilic-associated cellular damage.

## 2. Materials and Methods

### 2.1. Experimental animals

This study was approved by the Institutional Animal Care and Use Committee of the Feinstein Institutes for Medical Research and performed in accordance with the National Institutes of Health *Guide For The Care and Use Of Laboratory Animals* and *ARRIVE* guidelines. Male Sprague-Dawley rats, 5 weeks old and weighing 100–125g at the beginning of the experiment, were used, and all rats were between 350–450g at the time of collection (Charles River Laboratories, Kingston, NY). Animals were housed in a temperature-controlled room (12-h light/dark cycle), in cages lined with Enrich-o’Cobbs bedding (The Anderson, Inc, Maumee, OH) and were given access to food and water *ad libitum*.

### 2.2. Experimental groups

A total of 63 animals were acquired from Charles River Laboratories (Kingston, NY), with 60 undergoing training into adulthood (**Supplementary Material 1**). Three animals were excluded based on their performance and behavior during diving training to ensure result consistency. Following training, the successfully trained animals were allocated into the acute CGRP (aCGRP, aC) group, receiving a single session of intermittent DR (N=12); the chronic CGRP (cCGRP, cC) group, subjected to daily intermittent DR sessions over four weeks (N=12); the aCGRP group receiving the CGRP antagonist Fremanezumab (A-aCGRP, A-aC) (N=12); and the cCGRP group receiving the CGRP antagonist Fremanezumab (A-cCGRP, A-cC) (N=12). The remaining animals (N=12) were designated as the Sham (S) group for comparison. Half of each group underwent fresh sample collection and processing, while the other half underwent transcardial perfusion.

### 2.3. Rat voluntary diving protocol

Rats were trained to voluntarily dive through a multi-channel underwater tunnel using an adapted version of a previously validated method [42–44]. Rats were trained within a rectangular tank (100 x 38 x 15 cm) comprised of ¾-in thick acrylic pieces, with a ½-in thick top (MazeEngineers, Skokie, IL). The tank was modified to include 3 channels instead of the 5 channels specified in the original protocol. During a one-week period, the rats were acclimated to the aquatic environment and trained to voluntarily swim through the three-channel route, covering a total distance of approximately 2.5 meters. The training protocol involved initially acclimating the rats to the water environment to ensure their comfort and familiarity, followed by a gradual increase in the distance they were required to swim, ultimately covering the total 2.5 meters at a depth of approximately 10 cm in water maintained at 32±2°C. Rats were trained daily, with 3-5 trials per day per rat. Each rat was placed in the water at increasing distances from the finish platform. When the rats showed consistent successful negotiation of the path, they started on dive training to last another 2-weeks. It consisted of first developing the rats’ ability to dive underwater comfortably and voluntarily, and then gradually increasing the length of the underwater portion of the swimming path, until they showed consistent negotiation of the underwater multi-channel tunnel. Rats were required to show swimming ease within the first two days of training, voluntarily initiate dives within the first week of diving training and display minimal signs of agitation when handled or diving. Upon successful completion of training, rats in the acute CGRP group were subjected to a 30-minute diving session 3 days later, involving repeated dives every 5 minutes for a total of 7 dives, followed by sample collection (**Supplementary Material 1**). In contrast, rats in the chronic CGRP group participated in 30-minute repetitive diving trials 5 times per week over a 4-week period. On Day 50, the rats underwent a final 30-minute diving session followed by sample collection. Rats in the acute anti-CGRP group (N=12) and chronic anti-CGRP group (N=12) were administered Fremanezumab (Ajovy, 30 mg/kg, i.n., Teva Pharmaceuticals Industries Ltd., Parsippany, NJ) to inhibit CGRP. The acute anti-CGRP group received the treatment 5 hours before the final session and histological collection, while the chronic anti-CGRP group received Fremanezumab weekly leading up to Day 52. Sham rats, who underwent swim and diving training, were collected after a 3-day interval but did not participate in repetitive diving sessions.

### 2.4. Fresh Sample Collection and Processing

At the time of final sample collection, the animals were deeply anesthetized with isoflurane, and blood samples were obtained via the vena cava (N=6/group). The blood samples were refrigerated at 4°C for 30 minutes and subsequently centrifuged at 2000 × g for 10 minutes to separate the serum. The same animals were then decapitated, and tissue samples from the brain, lung, and kidney were collected. These tissue samples were rapidly frozen in liquid nitrogen and stored at −80°C until further analysis.

### 2.5. Transcardial Perfusion and Processing

A separate cohort of animals (N=6/group) was deeply anesthetized and subjected to transcardial perfusion with 0.1M ice-cold phosphate-buffered saline (Sigma, St. Louis, MO), followed by 4% paraformaldehyde. After perfusion, the heads were removed using a guillotine, and the brains, lungs, and kidneys were collected. The samples were fixed in 4% paraformaldehyde (Santa Cruz Biotechnology, Dallas, TX) overnight, then immersed in graded sucrose solutions, and cryoprotected in a mixture of 30% sucrose and optimal cutting temperature (OCT) compound (Electron Microscopy Sciences, Hatfield, PA). They were subsequently stored at −80°C until sectioning. The brains, lungs and kidneys were coronally sectioned at a thickness of 14 µm using a cryostat (Leica Biosystems, Deer Park, IL), mounted on Polysine glass slides (Thermo Fisher Scientific, Waltham, MA), and stored at −80°C until staining. Separate lung and kidney specimens were fixed in 4% formalin solution (Thermo Fisher Scientific, Waltham, MA) for histomorphological examination. Tissue processing, including paraffin embedding, sectioning at 4 µm thickness, and hematoxylin and eosin staining, was performed by AML Laboratories (St. Augustine, Florida).

### 2.6. Proteomics analysis

Aliquots of pulverized brain tissue from sham (N=4), aCGRP (N=4) and cCGRP (N=4) animals underwent comprehensive proteomic analysis via liquid chromatography-tandem mass spectrometry at Creative Proteomics (Shirley, NY, USA). Brain tissue was selected for analysis due to its relatively high constitutive CGRP expression and previously documented CGRP upregulation following trigeminal nerve activation, which occurs during DR [55–58]. Raw spectral data underwent standardized preprocessing including peak detection, peptide identification, and protein quantification. The resulting data was analyzed using a multi-platform bioinformatics workflow integrating R (MaxQuant v2.6.7.0, XLSTAT (v2024.4.2), and Python (Matplotlib v3.10.0) analytical environments. Data was subjected to median normalization followed by log-2 transformation. Missing intensity values were addressed through a hierarchical imputation strategy based on data sparsity patterns: k-Nearest Neighbor (kNN) for proteins with moderate missingness (n=1 missing values), Quantile Regression Imputation for proteins with substantial missingness (n=2 or 3 missing values), and Minimum Value Imputation for proteins with complete absence in one experimental group (n=4 missing values). Differentially expressed proteins were identified using dual selection criteria comprising Log2 fold-change thresholds >1.0 or <-1.0 combined with statistical significance (Student’s t-test with Benjamini-Hochberg correction, p<0.05). Visual representation of the proteomic landscape was accomplished through volcano plots and clustered heat maps generated using Python Matplotlib.

### 2.7. Western Blotting

Tissue samples from the brain, lung, and kidney were pulverized and aliquoted for subsequent analyses. Nuclear and cytosolic NRF2 levels were quantified using a commercial kit for nuclear and cytosolic protein isolation (**Supplementary Table 1**). Briefly, tissues were homogenized in a cytosolic pre-extraction buffer, followed by centrifugation. The resulting pellet was further homogenized in a secondary cytosolic extraction buffer and centrifuged, with the supernatant collected as the cytosolic extract. The remaining pellet was resuspended in nuclear extraction buffer, subjected to serial vortexing, and centrifuged, yielding the nuclear fraction. Total protein concentrations were determined in nuclear and cytosolic protein fractions, as well as in whole tissue homogenates prepared using RIPA lysis buffer (Thermo Fisher Scientific, Waltham, MA), supplemented with a protease and phosphatase inhibitor cocktail (Thermo Fisher Scientific, Waltham, MA). The lysates were resolved by SDS-PAGE, transferred to polyvinylidene difluoride (PVDF) membranes, and blocked with either 5% skim milk (Thermo Fisher Scientific, Waltham, MA) or 5% bovine serum albumin (Millipore Sigma, Burlington, MA) for 1 hour at room temperature. Membranes were then incubated overnight at 4°C with primary antibodies (**Supplementary Table 2**), which have been validated across various species and experimental conditions. Subsequently, membranes were incubated with HRP-conjugated secondary antibodies (Abcam, Waltham, MA) in 5% milk at room temperature for 1 hour. Protein signals were detected using chemiluminescence with ECL substrate (Thermo Fisher Scientific, Waltham, MA) on a Bio-Rad ChemiDoc Imaging System. ImageJ software was employed to quantify protein levels, and any adjustments to brightness or contrast were uniformly applied across the entire blot. β-actin and Lamin B1 served as loading controls for total and nuclear proteins, respectively. Western blot data are presented as fold change relative to sham controls to enable comparison across multiple blots. For sequesosome 1 (p62), adenosine monophosphate-activated protein kinase (AMPK), and sirtuin 1 (SIRT1), phosphorylated:total expression ratios were constructed using the β-actin corrected expressions, not the fold-change relative to sham.

### 2.8. GSH/GSSG Quantification

Levels of reduced (GSH) and oxidized (GSSG) glutathione were measured in powdered tissue samples from the brain, lung, and kidney, as well as in serum samples. Quantification of GSH, GSSG, and the GSH/GSSG ratio was performed using a commercial assay kit (**Supplementary Table 1**). Tissue concentrations of GSH and GSSG are reported in µmol/g, while serum concentrations are reported in µmol/mL.

### 2.9. Measurement of Malondialdehyde Levels

Lipid peroxidation was assessed in powdered tissue samples from the brain, lung, and kidney, as well as in serum samples, to evaluate free radical generation and oxidative stress. Malondialdehyde levels were quantified using a commercial assay kit (**Supplementary Table 1**), which measures thiobarbituric acid reactive substances as an indicator of malondialdehyde. Tissue samples were processed according to protocols optimized for high protein content.

### 2.10. RT-PCR

Total RNA was extracted from powdered brain, lung, and kidney tissues using Trizol reagent (Life Technologies, Carlsbad, CA). cDNA was synthesized from the isolated RNA using the High-Capacity cDNA Reverse Transcription Kit (Applied Biosystems, Foster City, CA). Quantitative RT-PCR was conducted to assess gene expression levels for heme oxygenase-1 (HO-1), superoxide dismutase (SOD), and NAD(P)H quinone dehydrogenase 1 (NQO1) (**Supplementary Table 3**). Primers were sourced from Eurofins Genomics (Louisville, KY). qPCR was performed using a 7500 Real-Time PCR System (Applied Biosystems, Foster City, CA) with SYBR Green PCR Master Mix reagents (Applied Biosystems, Foster City, CA). Rat GAPDH served as the endogenous control. Fold changes in gene expression were calculated using the delta-delta Ct method relative to control samples.

### 2.11. Immunofluorescence Staining and Quantification Analysis

Cryosectioned organ tissue slides were initially blocked with 5% goat serum (Abcam, Waltham, MA) in 1% bovine serum albumin (Sigma, Burlington, MA) for 1 hour at room temperature. The slides were then incubated overnight at 4°C with the primary antibody against either CGR, 4-HNE, or nitrotyrosine (**Supplementary Table 2**), followed by a 1-hour room temperature incubation with the corresponding secondary antibody. Subsequently, the slides were co-stained with anti-phosphorylated NRF2 antibody and its secondary antibody for 1 hour at room temperature. Slides were counterstained with DAPI and mounted using Vectashield Antifade mounting medium (Vector Laboratories, Burlingame, CA). Imaging was performed using the EVOS M7000 imaging system (Thermo Fisher Scientific, Waltham, MA) with a 20X objective and automated XY-stitching function to capture whole brain images. Signal expression was quantified semi-automatically in ImageJ [59,60], with pixel intensities ranging from 0 to 255, indicating the weakest and strongest intensities, respectively. Immunofluorescence data are presented as reciprocal intensity (arbitrary units, AU).

### 2.12. Assessment of Organ Morphological Alterations and Damage

Hematoxylin and eosin-stained slides were imaged using the EVOS M7000 imaging system (Thermo Fisher Scientific, Waltham, MA) with a 20X objective and automated XY-stitching function to capture whole lung and kidney images. Lung damage was assessed according to the degree of interstitial and pulmonary edema (0-2 each), as well as alveolar integrity (0-1) with a final score of 0-5 [61–63]. Kidney tubular injury, as defined by tubular atrophy, dilation, epithelial cell sloughing, and cast formation, as well as thickening of the tubular basement membrane was assessed, resulting in scores of 0-5 [64].

### 2.13. Statistical analysis

Data are presented as mean ± standard deviation (SD) with individual data points visualized in bar graphs. A priori power analysis with parameters α = 0.05 and β = 0.8 established a required minimum sample size of N = 6 per group. The Shapiro-Wilk and Kolmogorov-Smirnov tests were employed to assess the normality of distribution for all variables. Between-group comparisons (sham, acute CGRP, chronic CGRP, acute anti-CGRP, and chronic anti-CGRP) were performed using one-way analysis of variance (ANOVA) followed by Tukey’s post-hoc test for multiple comparisons. Statistical significance was defined as p < 0.05. All statistical analyses were executed using GraphPad Prism software (version 9, GraphPad Software, Boston, MA).

## 3. Results

### 3.1. Brain proteomics reveals NRF2-dependent antioxidant upregulation independent of oxidative stress markers

To investigate whether DR elevates NRF2-associated antioxidant proteins without concomitant increases in oxidative stress gene expression—a hallmark of non-electrophilic NRF2 activation—brain tissue specimens from both acute and chronic DR exposure paradigms were subjected to comprehensive proteomic analysis. Principal component analysis (PCA) was employed to assess inter-sample relationships and elucidate predominant sources of variance within the proteomic dataset (**Fig. 1A**). Notably, the chronic DR cohort demonstrated more pronounced segregation from sham controls compared to the acute DR group. The proteomic analysis encompassed 2388 distinct proteins, from which 102 were classified as antioxidative or oxidative stress-related entities, with particular emphasis on NRF2-modulated proteins. 35 proteins were identified as CGRP-associated proteins. Volcano plot representations delineate protein expression distribution in both acute and chronic exposure groups (**Fig. 1B**), with log2 fold change depicted on the abscissa and −log(p-value) on the ordinate. Statistical significance was defined by log2 fold change ≥1 or ≤-1 and P<0.5. The heat map visualization presents the most significantly dysregulated proteins, revealing an expression profile characteristic of NRF2 and CGRP pathway activation in the absence of electrophilic stimulation (**Fig. 1C**). Proteomic analysis revealed significant upregulation of NRF2-regulated antioxidant proteins, including *Gpx1, Gsta1/2/3/5, Gss, Glrx2, Txnrd2, Sod1, Prdx4,* and *Cox6a1,* which critically function in peroxide and superoxide detoxification pathways while enhancing glutathione synthesis [65–68]. In parallel, we observed significant downregulation of established oxidative stress markers, specifically *Adh7, Ogt,* and *Ndrg1* [69–71]. This tripartite reduction in markers suggests decreased retinol dehydrogenation— resulting in lower lipid peroxidation [72], diminished oxidative stress-induced protein glycosylation and oxidation [73], and reduced hypoxic signaling [74]—all of which are commonly promoted by classical electrophilic NRF2 activators [75,76]. Concurrently, the observed upregulation of antioxidant proteins, particularly those involved in glutathione synthesis and regulation, along with decreased oxidative stress biomarkers, provides compelling evidence for the non-electrophilic activation of the NRF2 pathway. Proteomic analyses further revealed significant upregulation of CGRP-associated proteins—including *Prkar2a, Prkar2b, Myh9,* and *Myl12b*—which act as downstream effectors in CGRP signaling cascades. Specifically, *Prkar2a* and *Prkar2b* modulate mechanisms that reduce oxidative stress by mediating L-citrulline and nitric oxide production and subsequently decreasing ROS levels, while *Myh9* and *Myl12b* are involved in the vasodilatory response characteristic of CGRP [77]. Notably, the chronic exposure cohort exhibited more pronounced alterations in protein expression compared to the acute exposure group across the majority of identified proteins. This observation aligns with the enhanced segregation revealed by principal component analysis and suggests a temporal dependency in the molecular response to endogenous CGRP activation.

**Figure 1:**
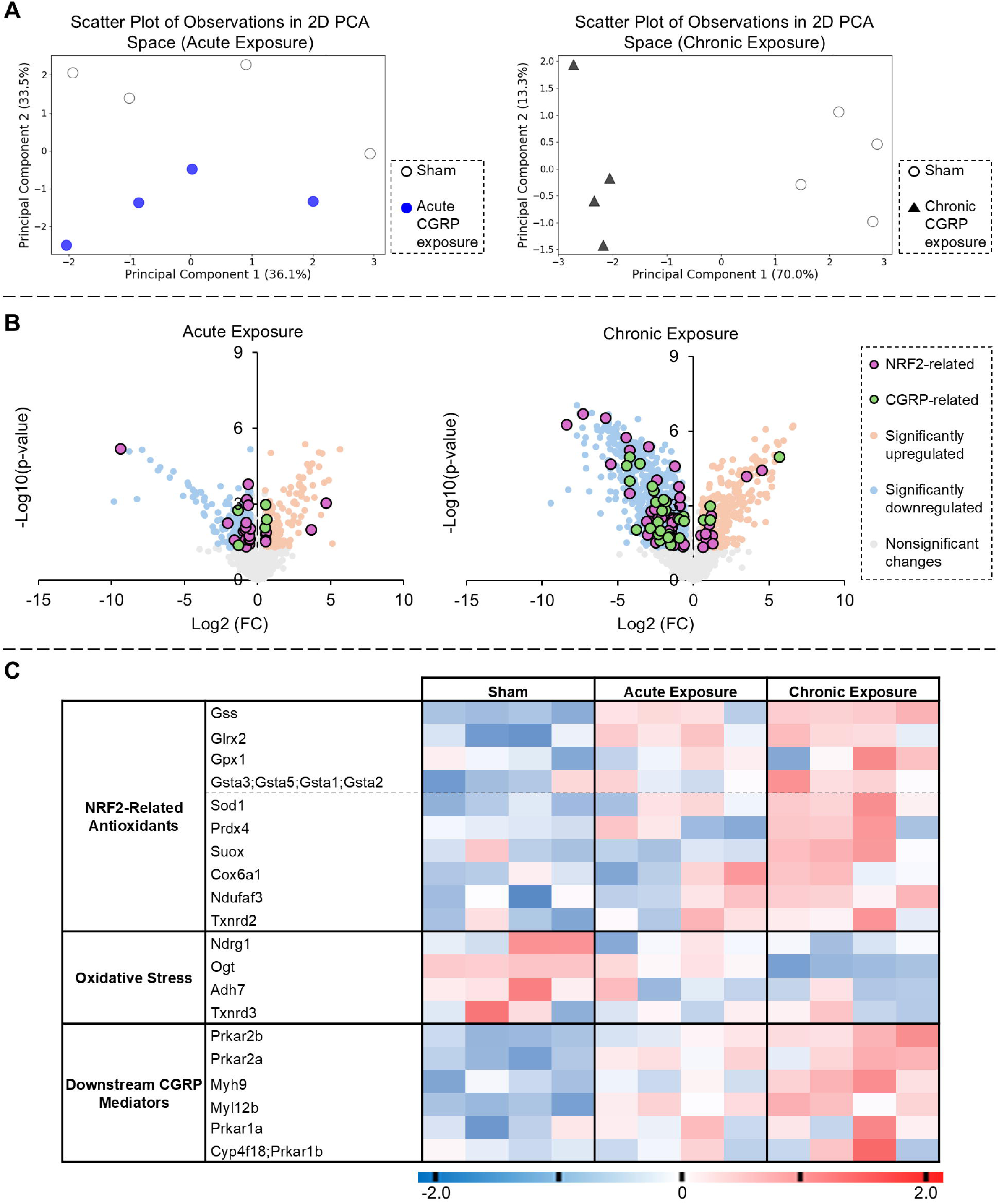
Brain proteomics reveals NRF2-dependent antioxidant upregulation independent of oxidative stress markers. **(A)** PCA analysis demonstrates distinct separation between sham and both acute and chronic CGRP exposure groups. **(B)** Volcano plot displaying fold change distribution of assessed proteins relative to sham (significance defined as fold change ≥2 or ≤0.5 and P<.05). **(C)** Heatmap showing differential expression of selected significantly modulated proteins categorized as oxidative stress markers, NRF2-mediated antioxidants, and CGRP pathway activation proteins. NRF2-mediated antioxidants are further subdivided by glutathione relation (above dotted line). (PCA: blue circles=sham, red triangles=CGRP exposure; Volcano plot: purple circles=NRF2-related, green circles=CGRP-related, red circles=significantly upregulated, blue circles=significantly downregulated, gray circles=nonsignificant changes).

### 3.2. CGRP enhances the translocation of NRF2 into the nucleus in both the brain and peripheral organs

To assess the extent of endogenous CGRP activation by DR, CGRP protein levels were measured in the brain, kidney, and lung following both acute and chronic DR exposures (**Fig. 2A-2D**). The brain exhibited the greatest increase in CGRP (∼3.5-fold, **Fig. 2B**), compared to ∼1.7-fold in the lung (**Fig. 2C**) and ∼2.2-fold in the kidney (**Fig. 2D**). No significant differences in CGRP expression were observed between acute and chronic exposures. Within the same animal, CGRP levels followed a trend of highest in the brain and lowest in the lung (**Supplementary** Fig. 2A-2B), indicating organ-specific variations in CGRP release from sensory nerves.

**Figure 2:**
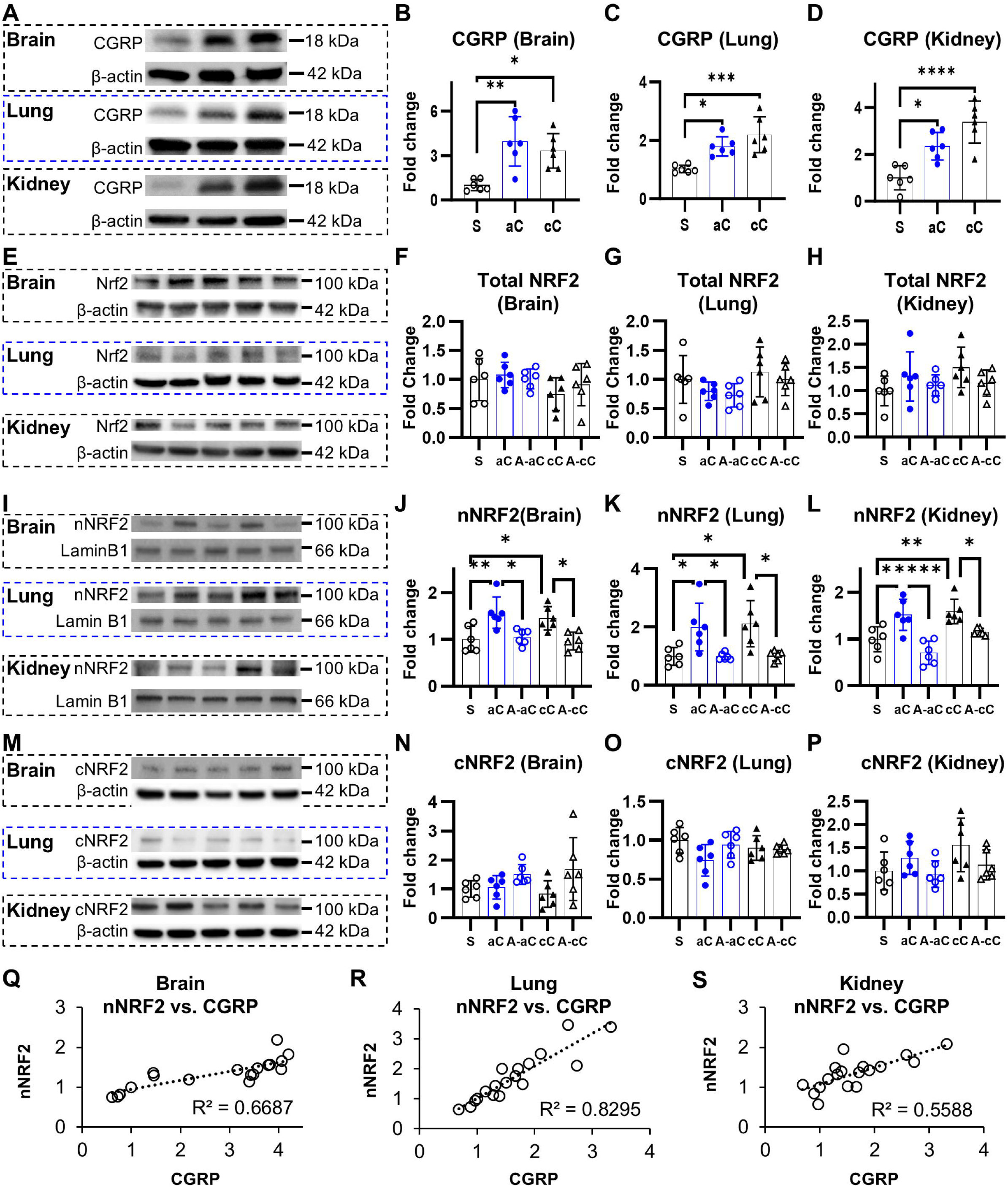
CGRP facilitates the nuclear translocation of NRF2 in both the brain and peripheral organs. **(A)** Representative Western blots illustrating CGRP expression levels. **(B-D)** CGRP expressions are significantly elevated in the brain, lung, and kidney during both acute and chronic DR; however, the differences between acute and chronic DR phases are not statistically significant. **(E)** Representative Western blots showing total NRF2 expression levels. **(F-H)** Among the studied organs, only the kidney exhibits an increase in total NRF2 levels in response to CGRP exposure. (I) Representative Western blots depicting nuclear NRF2 expression levels. **(J-L)** CGRP exposure enhances nuclear NRF2 expression in all three organs, an effect that is abolished by CGRP inhibition. **(M)** Representative Western blots demonstrating cytosolic NRF2 expression levels. **(N-P)** CGRP exposure does not lead to an increase in cytosolic NRF2 levels. **(Q-S)** CGRP expression is strongly correlated with nuclear NRF2 levels across all groups, with the lung displaying the strongest correlation and the kidney exhibiting the weakest. * *P* < 0.05, ** *P* < 0.01, *** *P* < 0.001, **** *P* < 0.0001 (S=sham; aC=acute CGRP; cC=chronic CGRP; A-aC=acute CGRP + Fremanezumab; A-cC= chronic CGRP + Fremanezumab).

Total, nuclear, and cytosolic NRF2 protein levels were quantified in the brain, kidney, and lung to evaluate NRF2 activation mediated by endogenous CGRP. In all three organs, total NRF2 levels did not exhibit significant changes (**Fig. 2E-2H**). However, nuclear NRF2 levels (**Fig. 2I-2L**) exhibited a significant increase in the brain (∼1.5-fold), lung (∼2-fold), and kidney (∼1.6-fold) in both acute and chronic CGRP-treated groups. Within individual animals, the most pronounced increase in nuclear NRF2 was observed in the lungs of both acute and chronic CGRP groups, whereas the kidney showed the smallest increase, suggesting organ-specific variations in response to CGRP (**Supplementary** Fig. 2C-2D). Despite the increase in nuclear NRF2, there was either no change, or a decrease, in cytosolic NRF2, with CGRP exposure (**Fig. 2M-2P**). The ratio of nuclear to cytosolic NRF2 expressions supported the activation of NRF2 (**Supplementary** Fig. 2E-2G), with marked increases in the brain (∼1.9-fold), lung (∼3-fold), and kidney (∼1.2-fold), indicating that the increase in nuclear translocation is not dependent on an increase in cytosolic NRF2 levels. Correlation analyses between nuclear NRF2 and CGRP levels (**Fig. 2Q-2S**) demonstrated strong positive correlations in the brain (R² = 0.6687) and lung (R² = 0.8295), and a moderate correlation in the kidney (R² = 0.5588), suggesting tissue-specific variations in CGRP-induced NRF2 activation.

A subset of animals treated with acute and chronic CGRP received Fremanezumab, a CGRP receptor antagonist, to validate CGRP as the primary mediator of NRF2 activation (**Fig. 2E-2P**). CGRP inhibition effectively blocked the DR-induced increase in nuclear NRF2 protein levels in the brain, lung, and kidney. Further, the inhibition of CGRP resulted in a decrease of the nuclear to cytoplasmic NRF2 ratio (**Supplementary** Fig. 2E-2G), contrasting directly to the increases observed without CGRP inhibition, and further supporting CGRP as the mediator of the observed NRF2 activation.

### 3.3. CGRP modulates NRF2 activity via phosphorylation in both the brain and peripheral organs

The impact of endogenous CGRP on promoting NRF2 phosphorylation at site s40 was further quantified. CGRP immunoreactivity was significantly elevated by DR in specific sub-regions of the brain, lung, and kidney (**Fig. 3A**, **Supplementary** Fig. 3A-3C). The most pronounced increases were observed in the cortex and CA1 regions of the brain, suggesting potential region-specific effects within the same organ. Post-hoc comparisons between the acute and chronic CGRP exposure groups revealed significantly higher CGRP expression in brain sub-regions such as the cortex, CA1, and corpus callosum in the chronic CGRP group, as compared to the acute group (**Fig. 3B**, **Fig. 3C**, **Supplementary** Fig. 3F). However, no differences were observed in overall protein levels, likely due to the use of homogenized tissue samples in protein measurements, which may mask region-specific variations.

**Figure 3:**
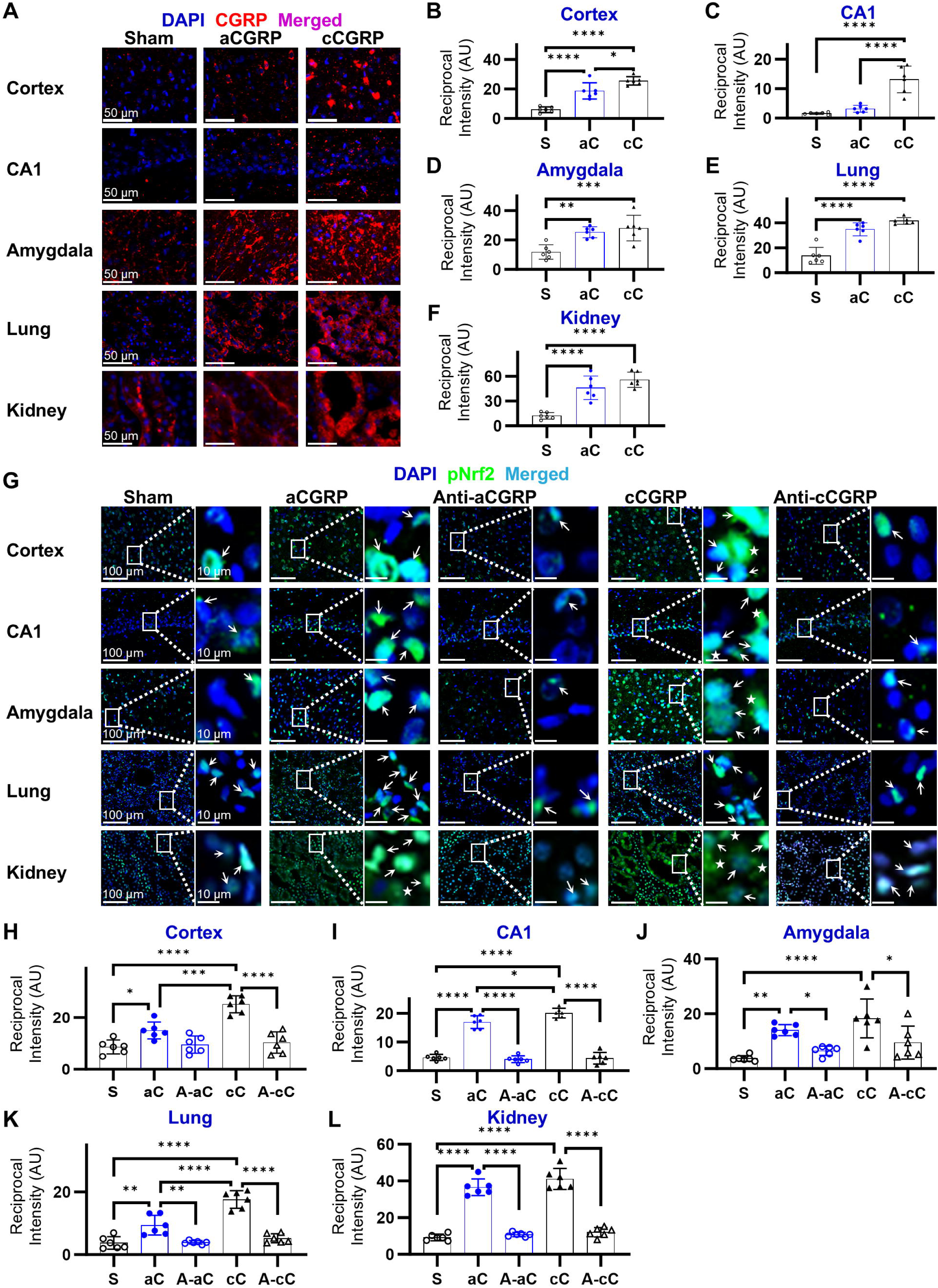
CGRP regulates NRF2 activity through phosphorylation in both the brain and peripheral organs. (**A-F**) Elevated expression of both intracellular and extracellular CGRP is observed in the dorsal cortex, CA1, amygdala, lung, and kidney during both acute and chronic DR. Chronic DR elicited a greater region-specific CGRP expression compared to acute DR. **(G-L)** Phosphorylated NRF2 expression is elevated in all brain subregions that exhibit increased CGRP levels, as well as in the lung and kidney. Chronic CGRP exposure significantly enhances phosphorylated NRF2 levels more than acute CGRP exposure in the cortex, CA1, and lung. Inhibition of CGRP negates the increase in phosphorylated NRF2 expression. Phosphorylated NRF2 expression is increased in the extranuclear space (depicted by lack of DAPI overlap) in the chronic cortex, CA1, and amygdala, as well as the kidney. * *P* < 0.05, ** *P* < 0.01, *** *P* < 0.001, **** *P* < 0.0001 (S=sham; aC=acute CGRP; cC=chronic CGRP; A-aC=acute CGRP + Fremanezumab; A-cC= chronic CGRP + Fremanezumab; arrow=examples of phosphorylated NRF2 and DAPI expression overlap; stars=regions where phosphorylated NRF2 does not overlap with DAPI).

CGRP exposure increased phosphorylated NRF2 expression in the brain, lung, and kidney (**Fig. 3G**, **Supplementary** Fig. 4A-4C). The relationship between CGRP and NRF2 phosphorylation was examined by measuring phosphorylated NRF2 intensity in regions with elevated CGRP. Both acute and chronic CGRP exposure led to significant increases in phosphorylated NRF2 in the dorsal cortex, CA1, amygdala, corpus callosum, dentate gyrus, and hypothalamus (**Fig. 3H-3J**, **Supplementary** Fig. 4D-4F). In the lung, phosphorylated NRF2 levels were elevated 3- to 5-fold following acute and chronic CGRP exposure, respectively, compared to Sham controls (**Fig. 3K**). In the kidney, these levels increased 4-fold with both acute and chronic CGRP exposure compared to Sham controls, with a visual increase in the amount of phosphorylated NRF2 which did not overlay with DAPI (**Fig. 3L**). Chronic CGRP exposure further elevated phosphorylated NRF2 expression in the brain and lung relative to acute exposure, although nuclear NRF2 levels remained unchanged. Chronic CGRP exposure also resulted in an increased appearance of phosphorylated NRF2 that did not co-localize with DAPI in the cortex, CA1, amygdala, and corpus callosum, suggesting its presence in the extranuclear space within these regions.

Inhibition of CGRP significantly reduced the previously observed increase in phosphorylated NRF2 within the brain, lung, and kidney sub-regions following both acute and chronic CGRP exposure (**Fig. 3G-3L**, **Supplementary** Fig. 4A-4C). Among the regions analyzed, the hypothalamus exhibited a notable increase in phosphorylated NRF2 after CGRP inhibition.

These findings suggest the presence of region-specific mechanisms underlying CGRP-dependent NRF2 activation, warranting further investigation.

Correlation analysis between phosphorylated NRF2 and CGRP expression revealed region-specific variations in correlation strength that were not discernible in whole organ homogenates (**Supplementary** Fig. 5). Notably, high correlation coefficients were observed in the cortex (R² = 0.8577), amygdala (R² = 0.858), and hypothalamus (R² = 0.7994). Other brain regions, such as the CA1 (R² = 0.5037), corpus callosum (R² = 0.4236), and dentate gyrus (R² = 0.5864), exhibited moderate correlations, suggesting a weaker influence of CGRP on NRF2 phosphorylation in these areas. In the lung (R² = 0.673) and kidney (R² = 0.7242), the correlations were relatively strong, further supporting CGRP as a mediator of NRF2 activation.

### 3.4. CGRP stimulates KEAP1-dependent pathways for NRF2 activation in the brain and peripheral organs

The expression levels of p62, phosphorylated p62 (ser349), and KEAP1 were assessed in the brain, lung, and kidney (**Fig. 4A-4C**) due to the known interaction between CGRP and p62 (Sequestosome 1) and the role of p62 in mediating KEAP1 in a non-electrophilic manner [78–80]. p62 protein levels increased in all examined organs, with the most significant elevation observed in the lung, where levels rose by 15-20% compared to the brain and kidney (**Fig. 4D-4F**). Phosphorylated p62 expression largely followed a similar trend across all three organs (**Fig. 4G-4I**). In contrast to p62 expression, KEAP1 protein expression decreased markedly in both acute and chronic CGRP exposure groups relative to Sham, with the most notable reduction occurring in the lung, especially in the acute exposure group (**Fig. 4J-4L**). Inhibition of CGRP almost completely reversed the changes in p62, phosphorylated p62, and KEAP1. Correlation analyses between p62, phosphorylated p62, and KEAP1 with nuclear NRF2 showed that p62 exhibited stronger correlations with nuclear NRF2 compared to KEAP1 (**Fig. 4M-4O**). The strongest correlations were observed in the lung (p62: R² = 0.6864, phosphorylated p62: R² = 0.7018, KEAP1: R² = 0.4939), aligning with the observed protein expression trends.

**Figure 4:**
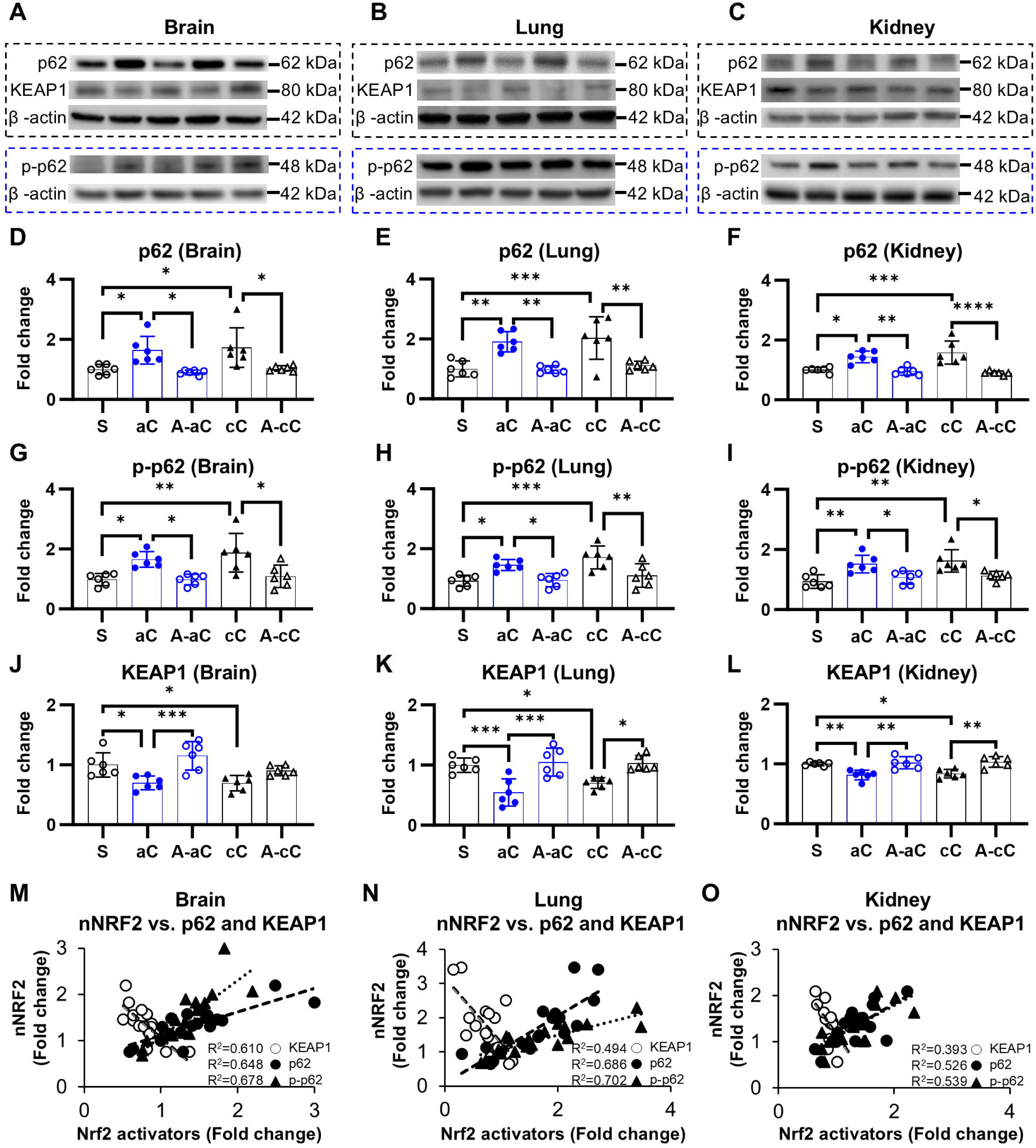
CGRP stimulates KEAP1-dependent pathways for NRF2 activation in the brain and peripheral organs. **(A-C)** Representative Western blots demonstrating the expression levels of p62, phosphorylated p62, and KEAP1. **(D-L)** p62 and phosphorylated p62 protein expression is significantly upregulated, while KEAP1 protein expression significantly decreases in the brain, lung, and kidney. Both alterations are abolished by CGRP inhibition. **(M-O)** Both p62 and KEAP1 exhibit fair to strong correlations with nuclear NRF2 across the brain, lung, and kidney, with p62 displaying the strongest correlations. * *P* < 0.05, ** *P* < 0.01, *** *P* < 0.001, **** *P* < 0.0001 (S=sham; aC=acute CGRP; cC=chronic CGRP; A-aC=acute CGRP + Fremanezumab; A-cC= chronic CGRP + Fremanezumab).

### 3.5. CGRP enhances KEAP1-independent, non-electrophilic pathways for NRF2 activation in the brain and peripheral organs

AMPK, phosphorylated AMPK (T183/T172), SIRT1, and phosphorylated SIRT1 (ser47), expression levels were evaluated in the brain, lung, and kidney due to their modulation by CGRP and the roles which AMPK and SIRT1 play in NRF2 activation [39,81] (**Fig. 5A-5C**). AMPK expression significantly increased in all examined organs (**Fig. 5D-5F**). AMPK levels in the brain following acute CGRP exposure were approximately two-fold higher than those observed in other groups or organs. Interestingly, though, phosphorylated AMPK levels only appeared to increase within the brain, suggesting an organ-specific effect (**Fig. 5G-5I**). SIRT1 and phosphorylated SIRT1 both increased to a comparable degree across all groups and organs (**Supplementary** Figure 6A**-6I**). Inhibition of CGRP nearly abolished the increases in AMPK, phosphorylated AMPK, SIRT1, and phosphorylated SIRT1. Correlation analyses revealed that AMPK, SIRT1, and phosphorylated SIRT1, had consistently high correlations with nuclear NRF2 protein levels (**Fig. 5J-5L, Supplementary** Figure 6J-6L).

**Figure 5:**
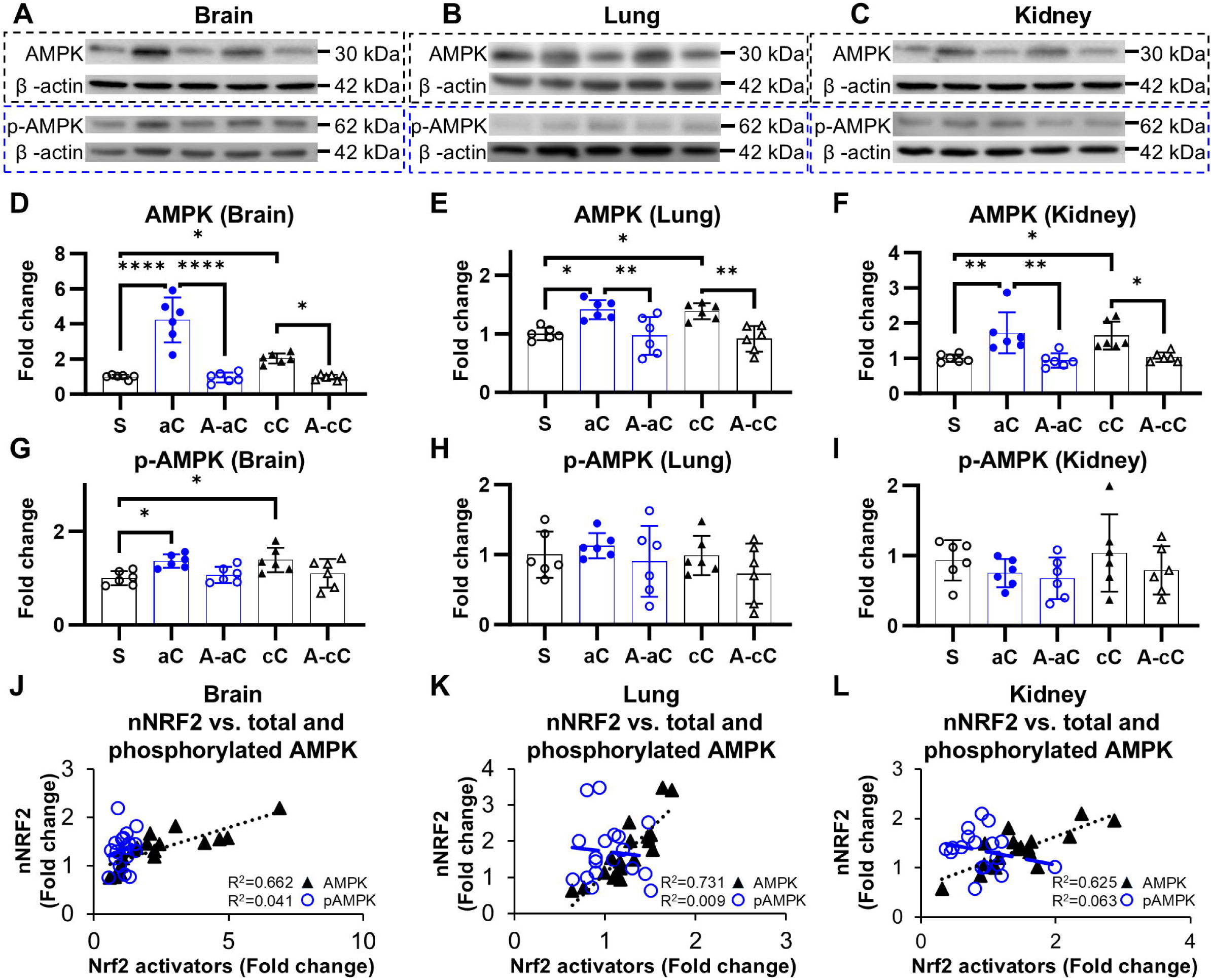
CGRP enhances KEAP1-independent, non-electrophilic pathways for NRF2 activation in the brain and peripheral organs. **(A-C)** Representative Western blots illustrating the expression levels of AMPK and phosphorylated AMPK. **(D-F)** AMPK protein expression levels are significantly upregulated in the brain, lung, and kidney following CGRP upregulation, with these changes being negated by CGRP inhibition. **(G-I)** Fold-change expression of phosphorylated AMPK does not change with CGRP exposure or inhibition, save for the chronic brain. **(J-L)** AMPK exhibits strong correlations with nuclear NRF2 in the brain, lung, and kidney. * *P* < 0.05, ** *P* < 0.01, *** *P* < 0.001, **** *P* < 0.0001 (S=sham; aC=acute CGRP; cC=chronic CGRP; A-aC=acute CGRP + Fremanezumab; A-cC= chronic CGRP + Fremanezumab).

### 3.6. CGRP enhances NRF2 activation in an organ-specific manner

The relative changes in the phosphorylated:total ratios of p62, AMPK, and SIRT1 were analyzed as a preliminary assessment of CGRP’s impact on NRF2 signaling activation (**Fig. 6**), evaluating the degree of activation relative to the baseline levels of each protein [82,83]. Notably, the marker showing the highest relative activation, suggesting the greatest impact by CGRP [82,83], exhibited organ-specific variation, with distinct differences observed between central (brain) and peripheral (lung and kidney) organs. While p62 showed the greatest degree of relative change in phosphorylation ratio within the lung and kidney, across both acute and chronic animals, AMPK exhibited the greatest degree of relative change within the brain. This was magnified in the chronic timepoint, with a ratio of ∼3.6 for AMPK, as compared to the p62 ratio of ∼1.4. Despite the increase in both total and phosphorylated SIRT1 levels, the calculated ratios, across all organs and times, were between 0.7-1.15, in direct contrast to either p62 or AMPK. As such, it is likely that SIRT1 does not play a dominant role in CGRP’s actions.

**Figure 6:**
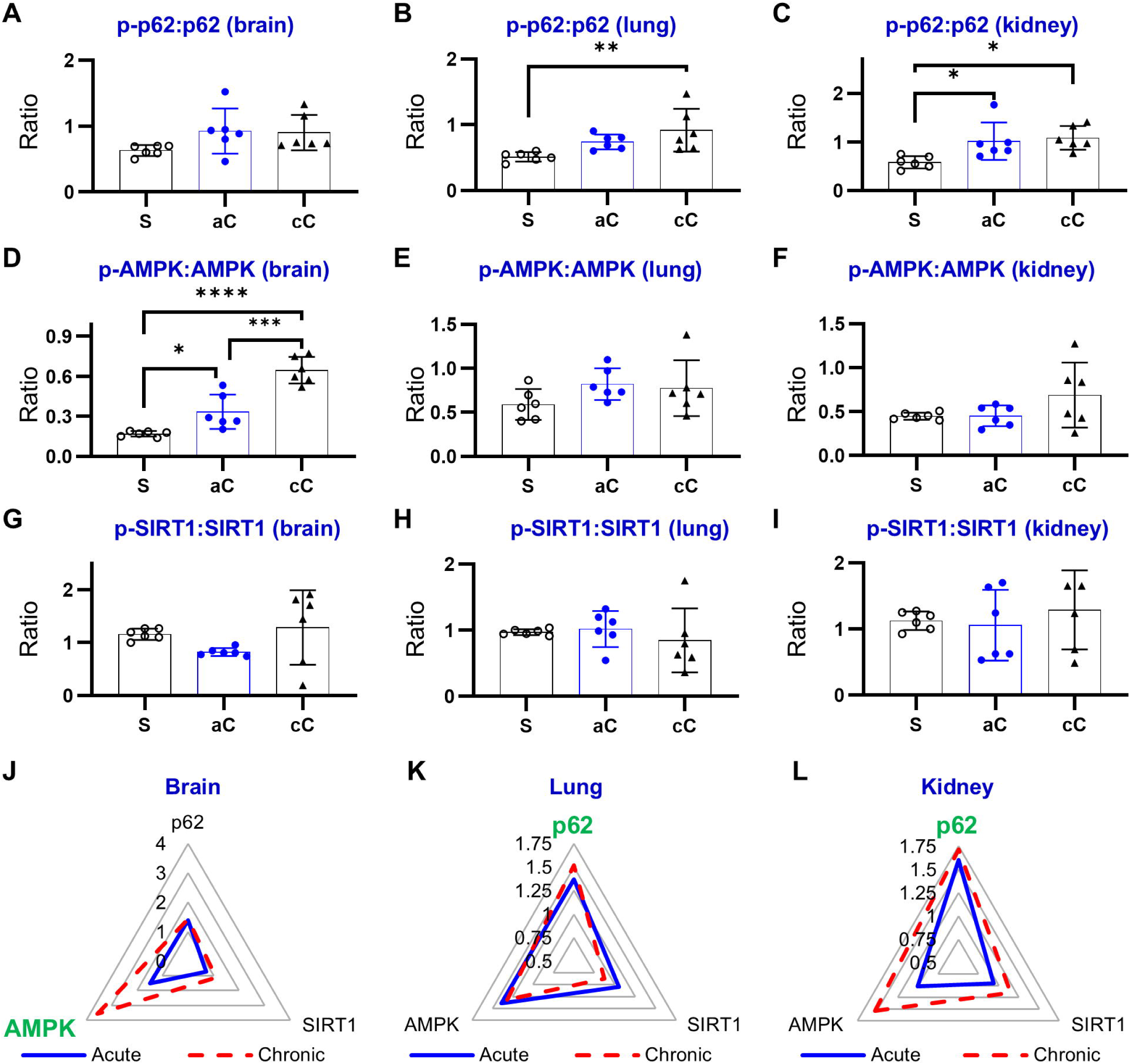
CGRP activates NRF2 signaling in an organ-specific manner. **(A-I)** The ratios of phosphorylated to total protein expression indicate that p62 is the most significantly affected mediator of NRF2 activation in peripheral organs (lung and kidney), whereas AMPK is the most impacted mediator in the brain. **(J-L)** Radar charts illustrating the relative changes in phosphorylation ratios across the three organs reveal that AMPK exhibits a more substantial change in the brain, whereas p62 shows a more pronounced change in the lung and kidney. (green label = identified most affected pathway)

### 3.7. CGRP stimulates potent transcription of NRF2-target antioxidant genes

mRNA and protein levels of NRF2 downstream genes, including HO-1, SOD, and NQO1, were measured in the brain, lung, and kidney (**Fig. 7**), to provide further confirmation of NRF2 activation [1–4,84]. In all three organs, acute and chronic CGRP exposure led to significant increases in PCR expression of all three antioxidant genes compared to sham conditions (**Fig. 7A-7I**). Notably, post-hoc analysis revealed that acute CGRP exposure resulted in significantly higher RNA expression levels of HO-1, SOD, and NQO1 in both the brain and lung compared to chronic exposure. Similarly, protein expression of HO-1, SOD, and NQO1 increased a significant degree with both acute and chronic exposure (**Fig. 7J-7U**).

**Figure 7:**
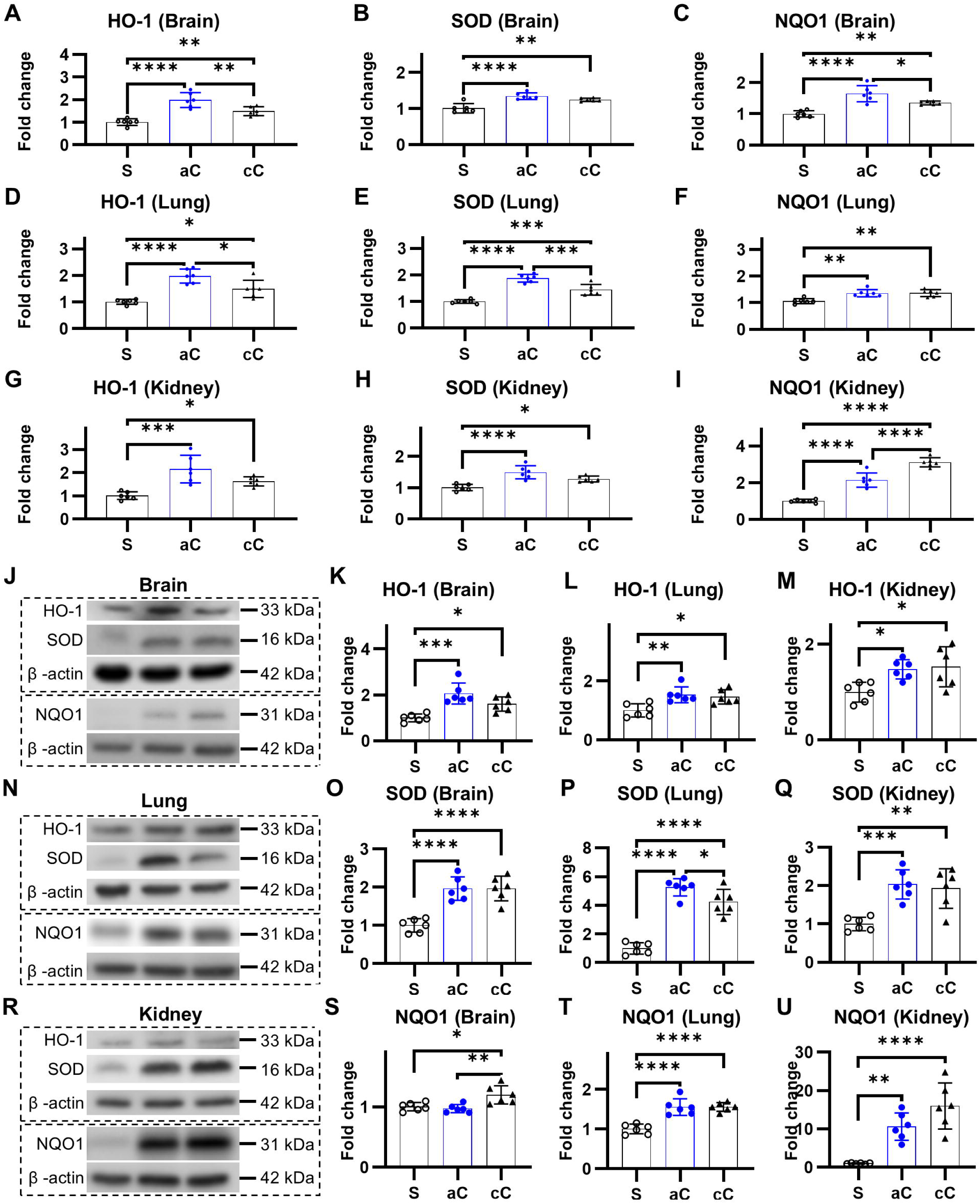
CGRP stimulates potent transcription of NRF2-target antioxidant genes. **(A)** Representative Western blots illustrating the expression levels of HO-1. **(B-D)** HO-1 protein expression levels are significantly upregulated in the brain, lung, and kidney following CGRP upregulation. **(E)** Representative Western blots illustrating the expression levels of SOD. **(F-H)** SOD protein expression levels are significantly upregulated in the brain, lung, and kidney following CGRP upregulation. SOD levels in the lung are more pronounced with acute exposure. **(I)** Representative Western blots illustrating the expression levels of NQO1. **(J-L)** NQO1 protein expression levels are significantly upregulated in the brain, lung, and kidney following CGRP upregulation. **(M-O)** Brain tissue exhibits significant elevations in the mRNA expression of HO-1, SOD, and NQO1. Notably, HO-1 and NQO1 levels are more pronounced with acute exposure. **(P-R)** The lung also exhibits significant increases in the mRNA expression of HO-1, SOD, and NQO1, with acute exposure resulting in greater elevations of HO-1 and SOD compared to chronic exposure. **(S-U)** The kidney shows significant elevations in the mRNA expression of HO-1, SOD, and NQO1. In contrast to the lung and brain, the expression levels of NQO1 in the kidney are higher following chronic exposure. * *P* < 0.05, ** *P* < 0.01, *** *P* < 0.001, **** *P* < 0.0001 (S=sham; aC=acute CGRP; cC=chronic CGRP).

### 3.8. CGRP activates NRF2 signaling in the absence of electrophilic indicators such as glutathione depletion, lipid peroxidation, and protein oxidation

CGRP’s molecular structure lacks functional groups, such as unsaturated carbonyl systems, that facilitate electrophilic reactions (**Fig. 8A**). To validate the absence of electrophilic mechanisms during endogenous CGRP-mediated NRF2 activation suggested by proteomic analysis, we quantified multiple oxidative stress biomarkers across serum, brain, lung, and kidney samples. Specifically, we measured: (1) glutathione levels, an indicator of electrophilic NRF2 activation corresponding to *Gss*, *Gsta*, *Glrx2,* and *Gpx* upregulation; (2) malondialdehyde and 4-hydroxynonenal concentrations, markers of lipid peroxidation correlating with *Adh7* downregulation; and (3) nitrotyrosine formation, a protein oxidation indicator associated with *Ogt* downregulation. GSH concentrations increased in the serum, lung, and kidney, but not in the brain (**Fig. 8B-8E**). The GSH/GSSG ratio increased across all samples (**Fig. 8F-8I**), indicating a lack of electrophilic activity. Furthermore, malondialdehyde levels decreased in the brain and kidney of chronic CGRP exposure animals, with no significant changes observed in other groups (**Fig. 8J-8M**). The lack of change in lipid peroxidation was confirmed via analysis of 4-HNE expressions in the brain, lung, and kidney (**Fig. 9A-9F, Supplementary** Figure 6A-6D). No increase was observed in any assessed region or organ, and, in fact, expression within the amygdala, hypothalamus, corpus callosum, and lung was significantly decreased. Assessment of protein oxidation, indicated by nitrotyrosine, indicated a concurrent significant decrease significantly in the cortex, CA1, amygdala, DG, and hypothalamus, and showed no change in the lung and kidney (**Fig. 9G-9L, Supplementary** Figure 6E-6H). The apparent lack, or decrease, of lipid peroxidation or protein oxidation increases indicates a lack of oxidative stress and further supports a non-electrophilic nature of endogenous CGRP’s NRF2 activation.

**Figure 8:**
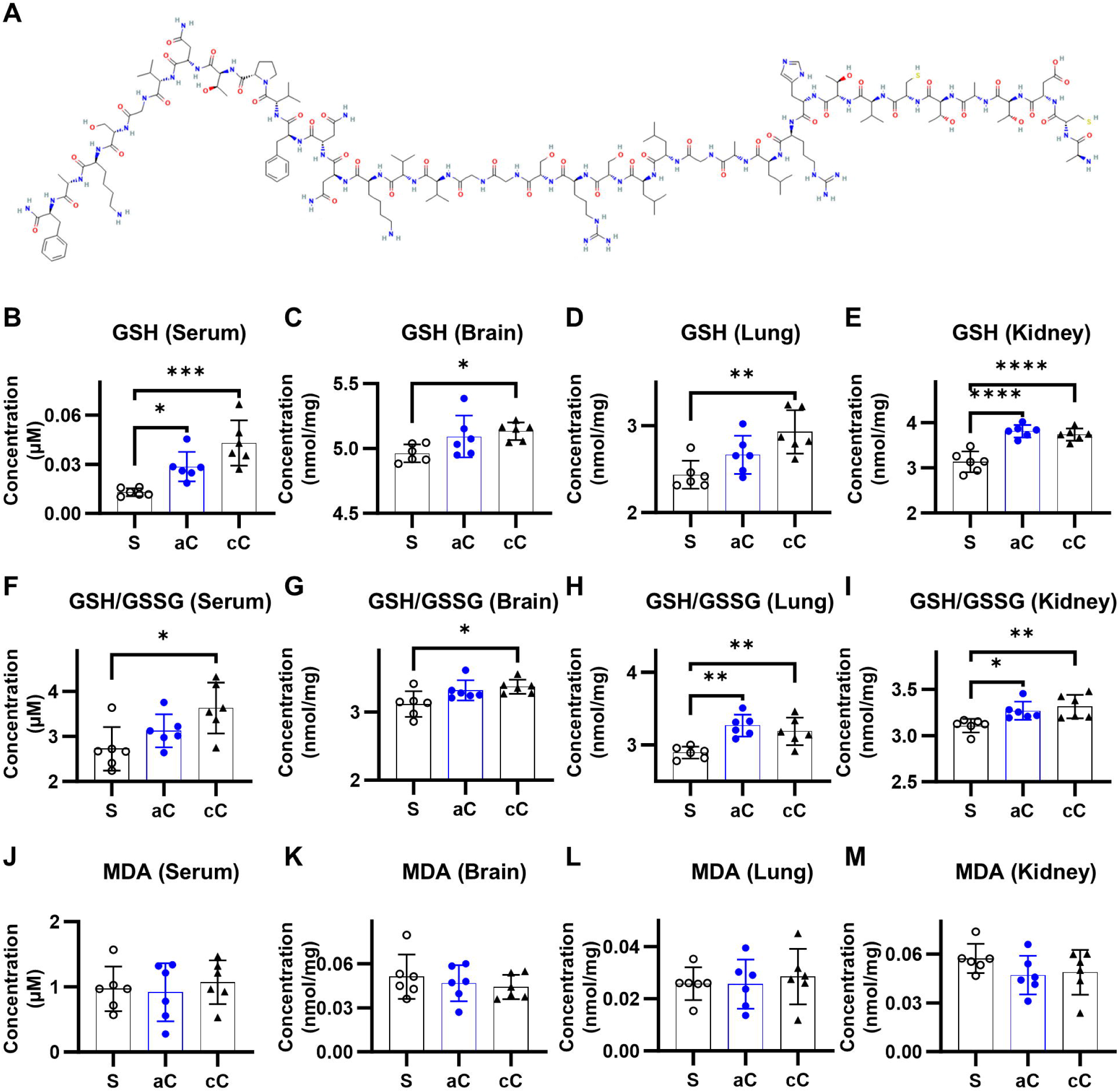
CGRP activates NRF2 signaling without triggering electrophilic stress, as indicated by the absence of glutathione depletion and increased lipid peroxidation. **(A)** The molecular structure of CGRP does not inherently promote electrophilic reactions. **(B–E)** CGRP enhances reduced glutathione (GSH) concentrations in the serum, brain, lung, and kidney, with statistically significant increases observed in the brain and lung only under chronic exposure. **(F–I)** CGRP exposure elevates the GSH/GSSG ratio across all studied tissues, with statistically significant increases detected in the serum and brain exclusively following chronic exposure. **(J–M)** Malondialdehyde levels exhibit a non-significant reduction in the brain and kidney, while serum and lung malondialdehyde levels remain unchanged. * *P* < 0.05, ** *P* < 0.01, *** *P* < 0.001, **** *P* < 0.0001 (S=sham; aC=acute CGRP; cC=chronic CGRP).

**Figure 9:**
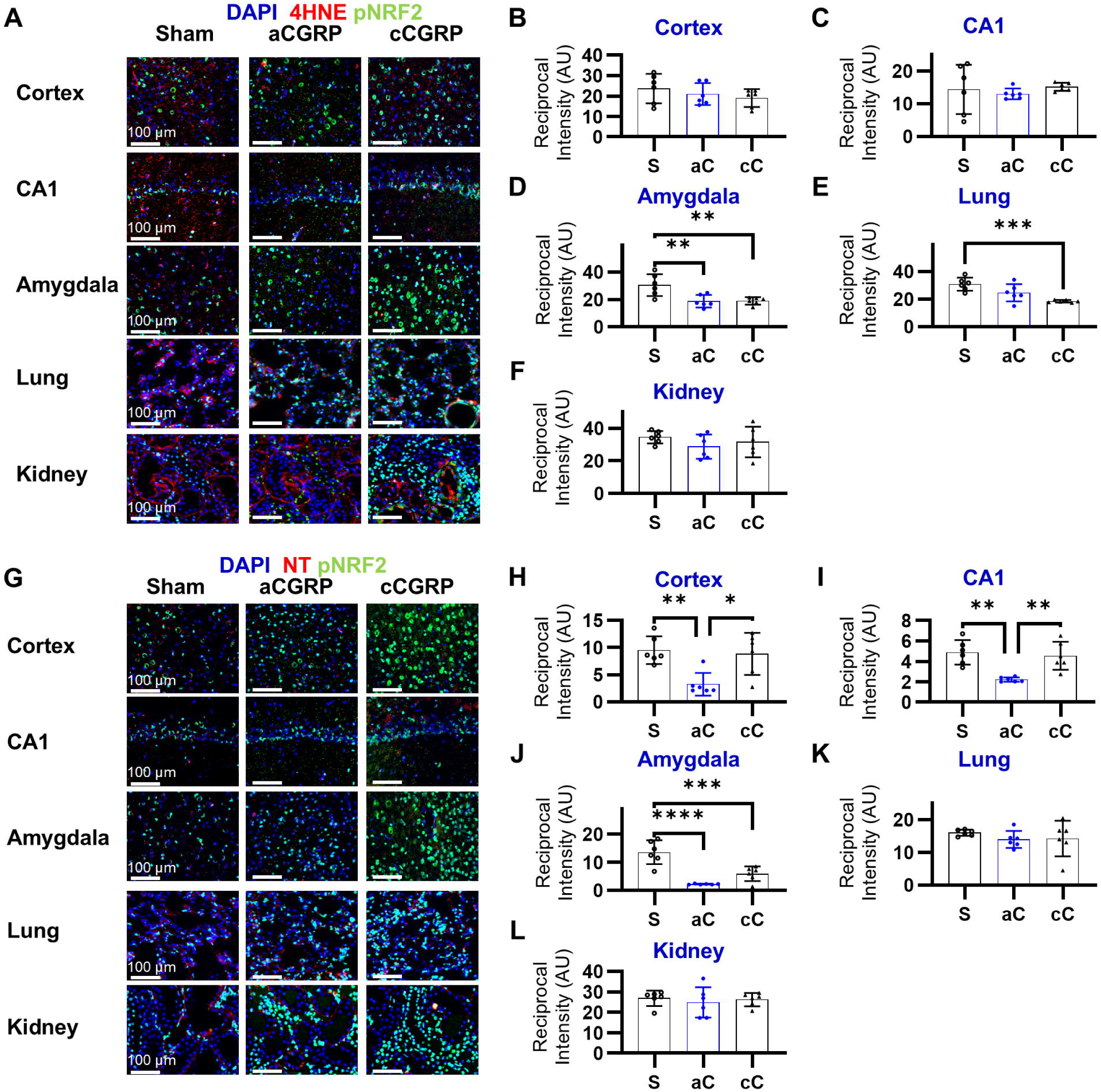
CGRP activates NRF2 signaling without inducing lipid peroxidation or protein oxidation. (**A–F**) Expression of 4-hydroxy-2-nonenal (4HNE) is reduced in the brain and lung, whereas levels in the kidney remain unchanged. **(G–R)** Nitrotyrosine expression is significantly decreased in the brain and lung, while it remains stable in the kidney. * *P* < 0.05, ** *P* < 0.01, *** *P* < 0.001, **** *P* < 0.0001 (S=sham).

### 3.9. Chronic CGRP-induced NRF2 activation does not lead to organ damage

Morphological damage within the lung and kidney was examined, as a means of comparing directly to the organ damage [5,12,85], in particular nephrotoxicity [85], previously observed with exogenous NRF2 activators. In the chronic CGRP exposure samples, no visual signs of nephrotoxicity, such as glomerular damage, kidney enlargement, or water retention were observed, nor was any indication of pulmonary damage (**Fig. 10**). Histological analysis of specimens from both acute and chronic exposure groups demonstrated morphology comparable to sham controls, suggesting favorable safety profile of endogenous CGRP as a physiological modulator of NRF2 signaling activation.

**Figure 10:**
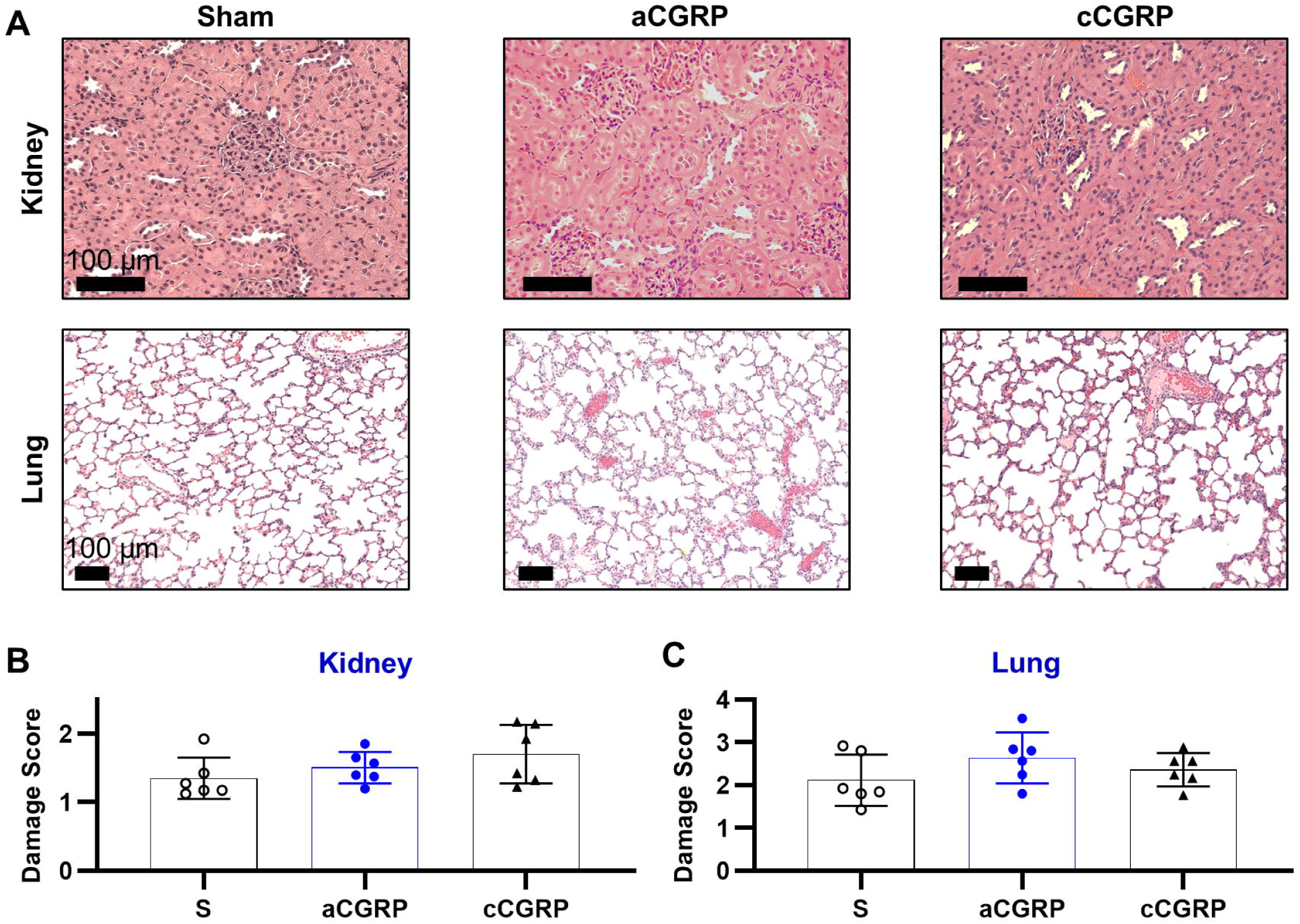
Chronic CGRP administration demonstrates absence of organ toxicity. **(A)** Representative micrographs displaying the morphology of kidney and lung tissues. **(B-C)** Histopathological analysis and damage grading of kidney and lung tissues reveal no evidence of structural damage after chronic CGRP exposure.

**Figure 11:**
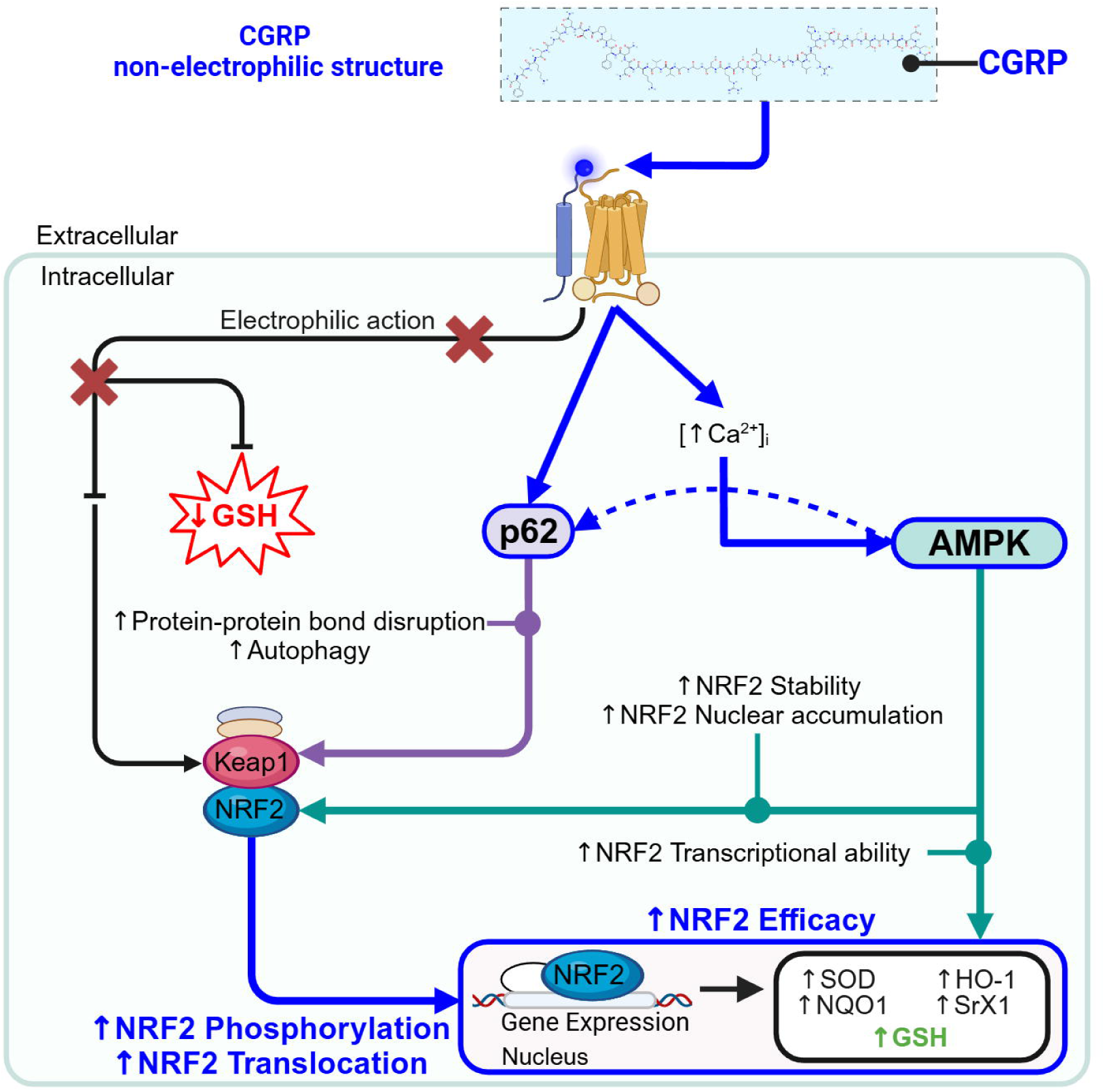
Mechanistic pathways of systemic CGRP-mediated NRF2 activation and redox homeostasis regulation. CGRP, secreted from sensory afferent terminals, exerts differential effects on central nervous system and peripheral tissues. The peptide’s molecular configuration precludes inherent electrophilic activity; instead, CGRP initiates NRF2 activation via AMPK- and p62-dependent signaling cascades, resulting in non-electrophilic NRF2 induction that maintains intracellular glutathione homeostasis. This signaling promotes dissociation of the KEAP1-NRF2 complex, consequently enhancing NRF2 stability, nuclear translocation, and DNA-binding activity. The downstream transcriptional program includes upregulation of phase II detoxifying enzymes and antioxidant molecules, with concurrent enhancement of functional antioxidant capacity and glutathione synthesis. Notably, tissue-specific mechanistic preferences exist within this pathway: peripheral tissues predominantly utilize p62-KEAP1-NRF2 signaling, while cerebral tissues preferentially engage AMPK-NRF2-mediated mechanisms, demonstrating organ-specific regulation of CGRP’s antioxidant effects. Abbreviations: CGRP, calcitonin gene-related peptide; GSH, glutathione; HO-1, heme oxygenase 1; KEAP1, Kelch-like ECH-associated protein 1; NRF2, nuclear related factor 2; NQO1, NAD(P)H quinone dehydrogenase; SOD, superoxide dismutase; SRX1, sulfiredoxin 1 (Created with BioRender.com).

## 4. Discussion

This study demonstrates that endogenous CGRP potently activates NRF2 signaling through non-electrophilic mechanisms in an organ-specific manner, with these effects being significantly attenuated by CGRP antagonist administration. CGRP-mediated NRF2 activation involves the p62-KEAP1-NRF2, AMPK-NRF2, and SIRT1-NRF2 pathways, with distinct regulatory dominance across tissues. Specifically, the KEAP1-dependent p62-KEAP1-NRF2 pathway predominates in peripheral organs, whereas the KEAP1-independent AMPK-NRF2 pathway is the primary driver of NRF2 activation in the brain (**Fig. 10**). Moreover, CGRP-induced NRF2 activation is associated with increased GSH levels, an elevated GSH/GSSG ratio, reduced protein oxidation and lipid peroxidation, and an absence of organ damage following chronic exposure, highlighting the non-electrophilic nature of this activation. The study further reveals organ-specific responses, including differential p62 expression across peripheral organs and distinct NQO1 expression patterns in response to CGRP. Additionally, acute CGRP exposure enhances antioxidant gene expression, whereas chronic exposure leads to increased phosphorylated NRF2 levels, indicating a time-dependent augmentation of oxidative stress resistance. Collectively, these findings indicate that CGRP primarily activates NRF2 through non-electrophilic pathways, offering a novel strategy for potent NRF2 activation while minimizing oxidative damage in conditions characterized by heightened cellular stress.

Endogenous CGRP upregulation, induced by DR, leads to significant NRF2 activation comparable to potent exogenous electrophilic activators like dimethyl fumarate (DMF) [86] and sulforaphane [87,88]. Proteomic profiling demonstrated significant upregulation of *Prdx4, Suox, Sod1, Glrx2,* and *Gss,* guiding subsequent analysis of NRF2 nuclear translocation and phosphorylation in brain and peripheral tissues. The CGRP-mediated NRF2 activation pathway exhibits mechanistic distinctions from DMF [75,76,86] and sulforaphane [87,88], characterized by non-electrophilic action as evidenced by elevated GSH levels and enhanced *Gpx1*, *Gsta*, and *Gss* expression, concomitant with reduced malondialdehyde, 4-HNE, nitrotyrosine, *Ogt*, *Adh7*, and *Txnrd3* expression. This non-electrophilic activation pattern likely stems from CGRP’s molecular structure, which lacks characteristic electrophilic moieties such as unsaturated carbonyl groups [89–93]. DMF, for example, contains an extended α,β-unsaturated carbonyl group that drives its high electrophilic reactivity, while CGRP consists of multiple nucleophilic zones with few unsaturated carbonyls [89], making it less prone to electrophilic interactions. Electrophilic NRF2 activators like DMF deplete glutathione by reacting with both reduced and oxidized forms, a hallmark of electrophilic action [7,8,10,94]. Within 60 minutes, DMF can significantly reduce detectable glutathione and form covalent glutathione conjugates, which may paradoxically elevate oxidative stress [10], a mechanism underlying its therapeutic use in psoriasis [95,96]. Only a limited number of pro-electrophilic drugs, which become electrophilic under oxidative stress, prevent glutathione depletion under normal conditions [7]. Due to the relative rarity of such compounds [8], non-electrophilic activators like CGRP offer promising alternatives for NRF2 activation without inducing glutathione depletion. Moreover, CGRP activates NRF2 much more rapidly than electrophilic activators, likely due to its reduced reliance on electrophile formation, with acute CGRP exposure resulting in NRF2 activation within 30 minutes, whereas DMF requires 8 to 24 hours to achieve similar effects [10,97]. This faster response emphasizes the potential of endogenous activators over exogenous ones. When compared to other endogenous NRF2 activators such as physical exercise, intermittent hypoxia, remote limb ischemic conditioning (RLIC), and VNS, CGRP shows advantageous properties as well. For instance, VNS elevates NRF2 in limited regions of the brain and does not exhibit non-electrophilic properties as CGRP does [20]. In contrast, physical exercise increases nuclear NRF2 levels by approximately 1.5-fold through ROS production [19,98], which, despite ROS’s electrophilic nature, does not lead to glutathione depletion but rather increases blood GSH levels [17]. Likewise, intermittent hypoxia and RLIC raise nuclear NRF2 by about 2-fold [17] and 1.6-fold [99], respectively, possibly via ROS release [100], which further differentiates CGRP’s mechanism. CGRP’s molecular structure, in combination with its ability to activate NRF2 through non-electrophilic pathways—characterized by increased GSH, reduced protein oxidation during both acute and chronic exposure, and decreased lipid peroxidation following prolonged administration—suggests its potential as a powerful NRF2 activator for therapeutic purposes.

The non-electrophilic activation of NRF2 by CGRP occurs in an organ-specific manner. In peripheral organs, the KEAP1-dependent p62-KEAP1-NRF2 pathway demonstrates stronger activation, whereas in the brain, the KEAP1-independent AMPK-NRF2 pathway is predominantly activated. These findings suggest that the CGRP-mediated activation of NRF2 is dependent on organ-specific regulatory mechanisms [82,83]. A significant upregulation of p62 and downregulation of KEAP1 were observed, accompanied by an increase in phosphorylated p62 levels. Notably, within the lung and kidney, p62 exhibited the highest phosphorylation ratios, suggesting a stronger CGRP-mediated impact and a predominant effect in these organs. p62 promotes KEAP1-dependent NRF2 activation by disrupting the KEAP1-NRF2 complex through protein-protein interactions [4,101], leading to the release of NRF2, enhanced nuclear translocation, and increased autophagic degradation of KEAP1 [102,103]. This process is further amplified by phosphorylation of p62 at serine 351 (serine 349 in rats), which alters the crystal structure of p62, thereby enhancing its binding affinity to KEAP1 [104,105]. While p62 plays a critical role in NRF2 activation by promoting the dissociation of NRF2 from the KEAP1-NRF2 complex [4,101], positioning this pathway as a primary upstream regulator and the most impacted by CGRP in certain contexts, this does not preclude the possibility that, under specific conditions— such as those in the brain—an alternative pathway may be more prominently influenced by CGRP. In the brain, the conditions preferentially support AMPK activation as the primary mechanism for KEAP1-independent NRF2 activation. AMPK facilitates this process by hyper-phosphorylating NRF2 at specific serine residues, leading to the upregulation of downstream target genes [106–108], thereby enhancing NRF2 nuclear accumulation, functional activity, and transcriptional potency [92–96]. The pronounced increase in AMPK phosphorylation within the brain, compared to peripheral organs, further potentiates NRF2 activation in this context [113,114]. However, while phosphorylation enhances AMPK activation, it is not an absolute requirement for its function [115].

Therefore, the absence of increased phosphorylated AMPK in peripheral organs does not necessarily indicate a lack of activation, whereas the presence of elevated phosphorylation in the brain suggests a significantly greater impact of CGRP on AMPK signaling. For comparison, sulforaphane has been shown to upregulate AMPK in the heart without inducing its phosphorylation, despite partially mediating NRF2 activation through AMPK [115]. This distinction may explain why sulforaphane administration under healthy conditions does not lead to increased NRF2 activation. The brain’s unique cellular makeup may affect the way in which AMPK functions. Both total AMPK and AMPK phosphorylation can be increased by intracellular calcium levels ([Ca^2+^]_i_) [116]. In the brain, CGRP is known to significantly elevate [Ca^2+^]_i_, owing to the high percentage of neuronal cells in this region [117–119]. Consequently, the brain may exhibit a preference for AMPK activation over p62, despite AMPK’s role in NRF2 activation existing after its release from the NRF2-KEAP1 complex. Collectively, these findings demonstrate that CGRP activates NRF2 in an organ-specific manner, with the p62-KEAP1-NRF2 pathway predominantly involved in the lung and kidney, while the AMPK-NRF2 pathway dominates in the brain.

Alongside the distinct primary pathways observed in the brain and peripheral organs, findings in the peripheral organs further support the notion of CGRP exerting organ-specific effects on NRF2 activation. Specifically, (1) the lung demonstrates heightened sensitivity to CGRP exposure, as indicated by particularly elevated nuclear NRF2 and p62 levels, and (2) the kidney exhibits unique expression patterns of NQO1. The lung shows the most pronounced increase in nuclear NRF2, accompanied by significant alterations in p62 and KEAP1, thereby reinforcing the concept of p62 being the predominant pathway in peripheral organs. Physiologically, CGRP application to areas with reduced sensory nerve innervation triggers a more substantial pain and sensitization response compared to more innervated regions [55–57]. While αCGRP is abundant in the lung, its relatively low innervation compared to the kidney or brain [28,120,121] may enhance the CGRP response, increasing p62 levels, promoting autophagic degradation of KEAP1, and further activating NRF2 [102,103,122,123]. Distinct effects are further observed in the kidney, which uniquely shows increased expression of the antioxidant gene NQO1 following chronic, rather than acute, CGRP exposure. NQO1 is involved in self-reinforcing feedback loops within the kidney [124,125], particularly under conditions of mitochondrial modification [15,126]. The kidney’s heightened sensitivity to mitochondrial bioenergetic changes, attributed to its high levels of long-chain polyunsaturated fatty acids and elevated mitochondrial content [127], predisposes it to these feedback mechanisms. These findings suggest that the organ-specific modulation of NRF2 signaling by CGRP is influenced by unique structural characteristics and sensitivity, warranting further investigation.

Our findings suggest that CGRP’s activation of NRF2 follows a time-dependent pattern, with the duration of exposure playing a critical role. Acute CGRP exposure results in significantly greater upregulation of NRF2-targeted antioxidative gene expression compared to chronic exposure, despite not expressing increased NRF2 activation. This may be indicative of a change in oxidative stress sensitivity following chronic exposure. Four weeks of CGRP exposure preserves nuclear NRF2 levels similar to acute exposure while further increasing phosphorylated NRF2 (∼11.0-fold vs. ∼6.3-fold). Although phosphorylated NRF2 typically translocates to the nucleus, some remains in the cytoplasm [128], which would not be picked up during examination of cytosolic NRF2 levels. CGRP’s enhancement of oxidative stress resistance [29,31,32,129,130] may decrease the sensitivity of nuclear import mechanisms, creating a disparity between the production and translocation rates of phosphorylated NRF2 [131]. Thus, stable nuclear NRF2 levels can be maintained as long as NRF2 phosphorylation continues to rise, even with reduced translocation rates. This is particularly apparent in the cortex, CA1, amygdala, and corpus callosum of the chronic exposure animals, where, in addition to an increase in the absolute levels of phosphorylated NRF2 and its overlaying the nuclei, there is also a notable increase in the expression of phosphorylated NRF2 within the extranuclear space. The decrease in nuclear import mechanics may also be indicative of a decrease in oxidative stress sensitivity, resulting in a relative decrease in antioxidant gene mRNA expression. Despite this decrease in antioxidative gene expression, however, the absolute changes in expression levels are comparable to those induced by exogenous electrophilic NRF2 activators such as DMF [132] and sulforaphane [87]. Temporal proteomic profile alterations further corroborate exposure duration-dependent modulation of CGRP- and NRF2-mediated signaling, evidenced by augmented expression patterns of *Ogt*, *Adh7*, *Sod1*, *Gst*, *Gpx*, *Gss*, and *Prkar2b* following chronic CGRP administration. Time-dependent effects are also supported by the expression patterns of AMPK. In addition to the observed shift in AMPK phosphorylation in the brain following chronic CGRP exposure, there is a shift in the total amount of AMPK protein expression, with only a 2X increase with chronic exposure, as compared to the 4X increase present with acute exposure. The aforementioned increase in [Ca^2+^]_i_ which likely occurs within the brain can cause an upregulation in total AMPK expression with short-term exposure [118,119,133], while prolonged exposure diminishes this effect [133]. However, the same increase in [Ca^2+^]_i_ increases the amount of AMPK phosphorylation [116], which likely increases the efficacy of the present AMPK, thus maintaining AMPK as a dominant pathway, despite a relative decrease in total expression. Altogether, this study may be the first to elucidate distinct temporal activation patterns of NRF2 signaling pathways in response to CGRP, highlighting the necessity for further investigation.

This study has certain limitations. Firstly, our data suggest differential CGRP-mediated NRF2 activation mechanisms across tissue types—p62-KEAP1-NRF2 signaling is predominant in peripheral organs (lung and kidney) versus AMPK-NRF2 pathway in brain tissue. However, these conclusions derive from indirect assessments without selective pathway inhibition. This methodological limitation was necessary as chronic inhibition of p62 or AMPK is associated with neurodegenerative and metabolic pathologies [134–136], and our primary objective was to evaluate physiological responses to chronic CGRP supplementation, precluding the use of selective inhibitors [137–139]. Furthermore, investigating organ-specificity necessitated whole-organ analysis rather than *in vitro* models to accurately characterize intercellular interactions. Secondly, exogenous CGRP administration was excluded as a comparative model due to its established hypotensive effects [140,141], limited blood-brain barrier penetration [142,143], and short serum half-life. Instead, we employed the DR to induce endogenous CGRP release, which offers several physiological advantages: concurrent peripheral vasoconstriction preventing hypotension [42–45,51,144], effective targeting of both central and peripheral tissues, and prolonged CGRP half-life with release into cerebrospinal and extracellular fluids [77,145]. DR is currently employed as a clinical treatment for supraventricular tachycardia, highlighting its potential as a method for enhancing endogenous CGRP levels [77]. Furthermore, precisely controlled electrical trigeminal nerve stimulation has been shown to biomimic DR, offering an alternative for patients unable to undergo traditional DR induction due to age or trauma, thus demonstrating its broad clinical applicability [58].

## 5. Conclusion

This study demonstrates that elevated endogenous CGRP, induced by DR, results in significant non-electrophilic activation of NRF2 in both the brain and peripheral organs. This effect is characterized by increased levels of nuclear and phosphorylated NRF2, enhanced nuclear translocation rates, upregulation of NRF2-targeted antioxidative gene transcription, and activation of non-electrophilic NRF2 pathways. Additionally, there is an increase in glutathione reserves, a reduction in lipid and protein oxidation, and a lack of organ damage with chronic exposure. These characteristics differentiate CGRP from traditional electrophilic NRF2 activators, suggesting that CGRP facilitates the concurrent derepression, stabilization, aggregation, and activation of NRF2, potentially enabling synergistic modulation of NRF2. The organ-specific nature of the most affected pathways, with the KEAP1-dependent p62-KEAP1-NRF2 pathway being predominantly affected in peripheral organs and the KEAP1-independent AMPK-NRF2 pathway being most impacted in the brain, underscores the need for further investigation. This necessity is further emphasized by the time-dependent patterns of NRF2 activation and its varying potency. These findings suggest that CGRP may serve as a therapeutic agent for modulating NRF2 pathways in both acute and chronic diseases while minimizing cellular stress by bypassing electrophilic reactions, particularly in conditions characterized by rapid and significant increases in oxidative and cellular stress.

## Supporting information

Whole Blot Images

## CRediT authorship contribution statement

**K. Powell**: Methodology; formal analysis; writing – original draft preparation; Writing – review & editing. **S. Wadolowski**: Methodology; formal analysis; writing – original draft preparation. **W. Tambo:** Methodology; formal analysis; writing – original draft preparation. **E. Chang:** Writing – review & editing. **D. Kim:** Methodology. **A. Jacob:** Writing – review & editing. **D. Sciubba:** Writing – review & editing. **Y. AlAbed:** Writing – review & editing. **P. Wang:** Writing – review & editing. **C. Li**: Conceptualization; investigation; formal analysis; methodology; writing – original draft preparation; writing – review & editing; funding acquisition; project administration; resources; supervision.

## Funding resources

This work is supported in part by the United States Army Medical Research Acquisition Activity (USAMRAA) under award #HT9425-24-1-1017, the US Army Medical Research and Materiel Command (USAMRMC) under award #W81XWH-18-1-0773, the Zoll Foundation Award, and the merit-based career enhancement award at the Feinstein Institutes for Medical Research.

## Acknowledgements

We would like to thank Dr. Pnina Powell for her aid in assessing the chemical structures of CGRP and DMF.

## Data Availability Statement

Individual data points are plotted for all graphs. The data that support the findings of this study are available from the corresponding author upon reasonable request.

## Ethics Approval Statement

Experiments within this study were approved by the Institutional Animal Care and Use Committee of the Feinstein Institutes for Medical Research and performed in accordance with the National Institutes of Health *Guidelines For The Use Of Experimental Animals*.

---

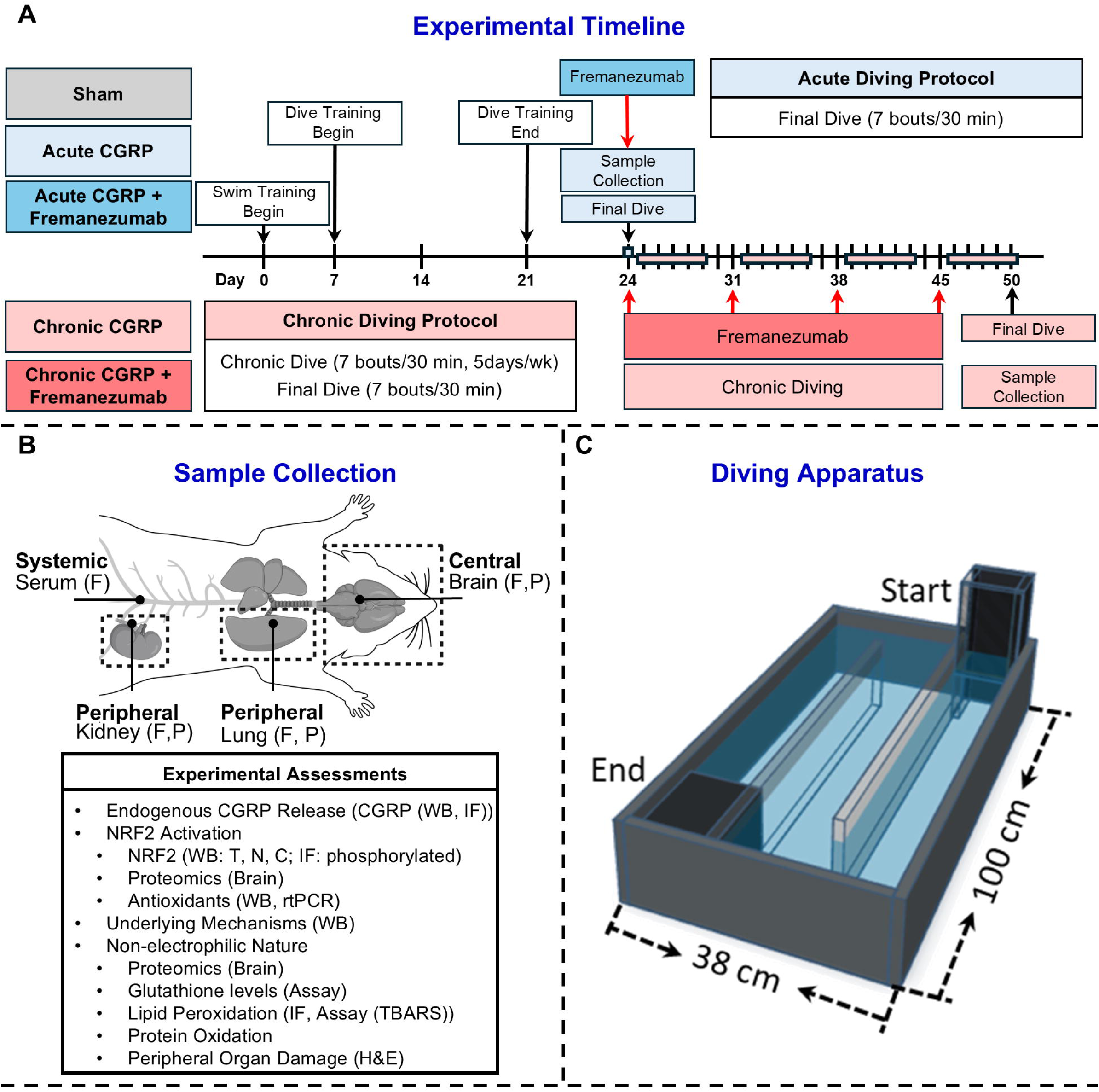

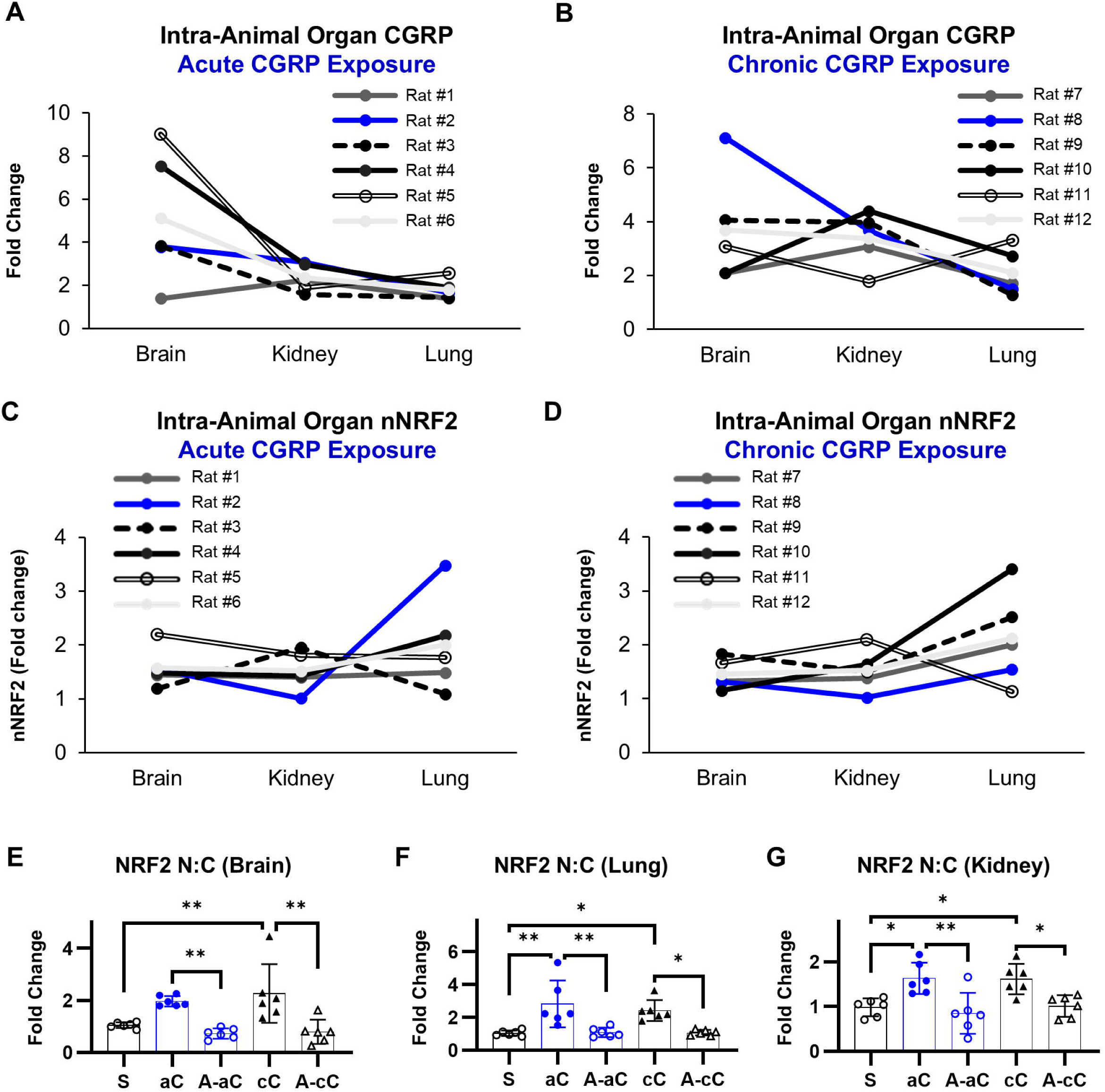

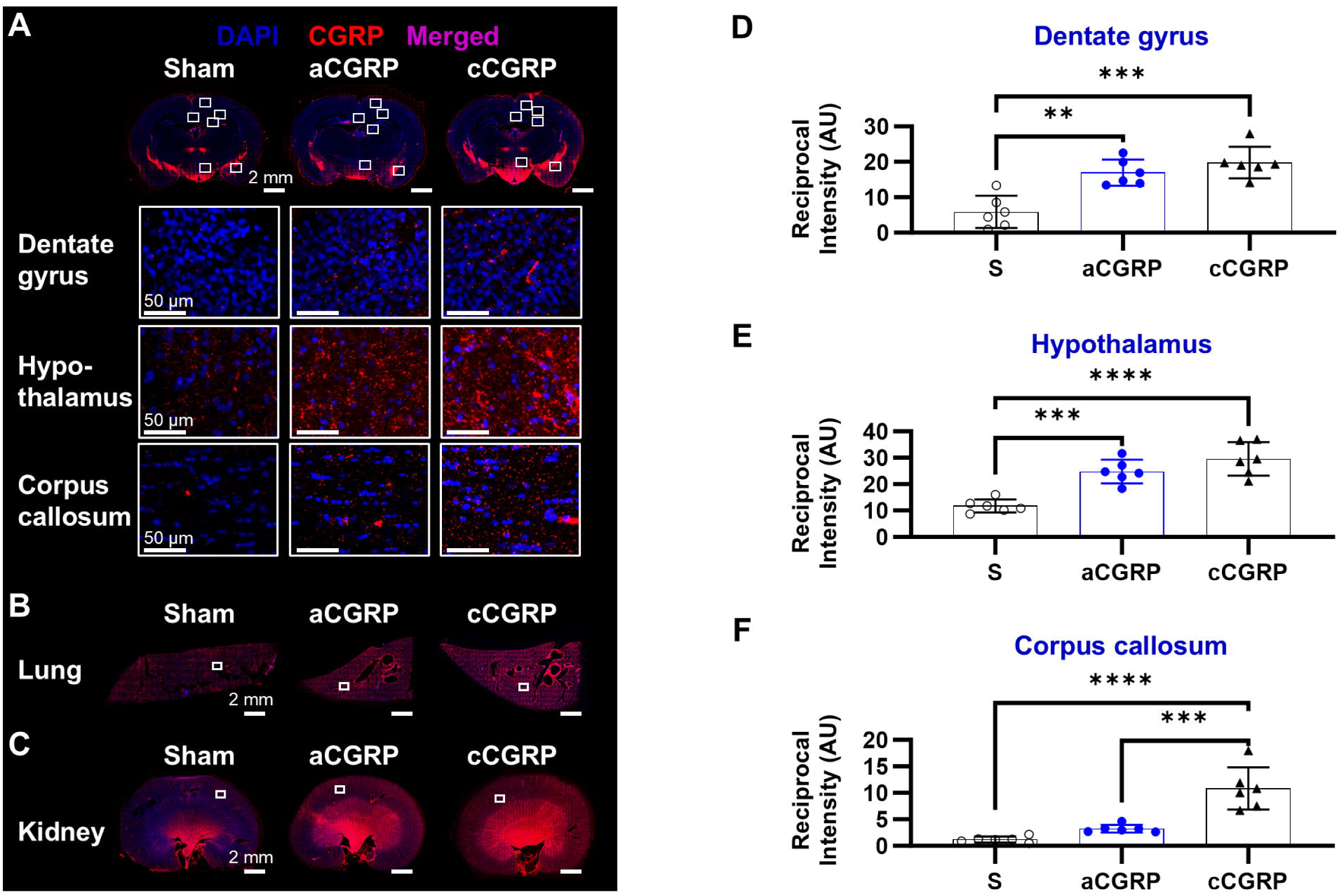

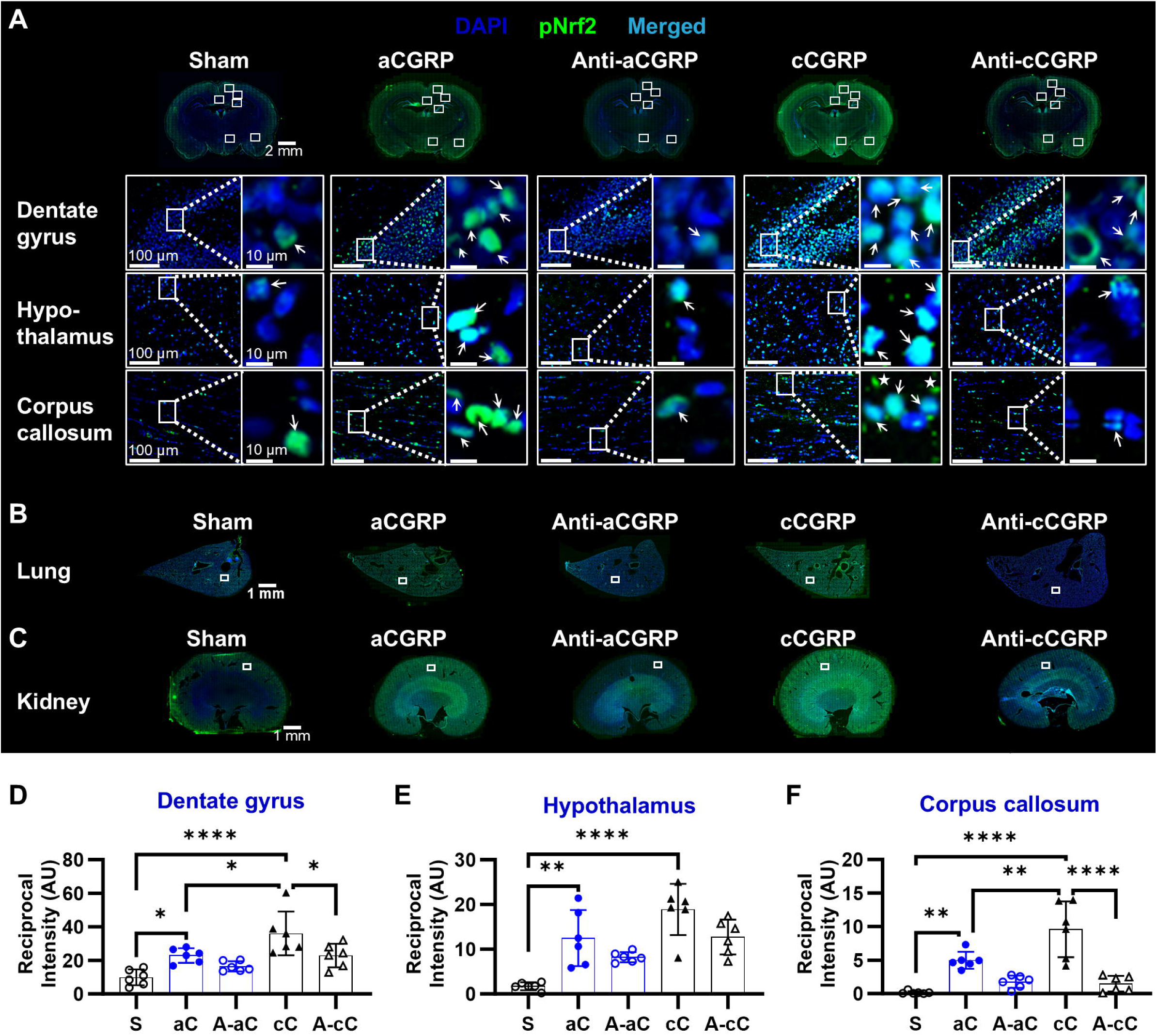

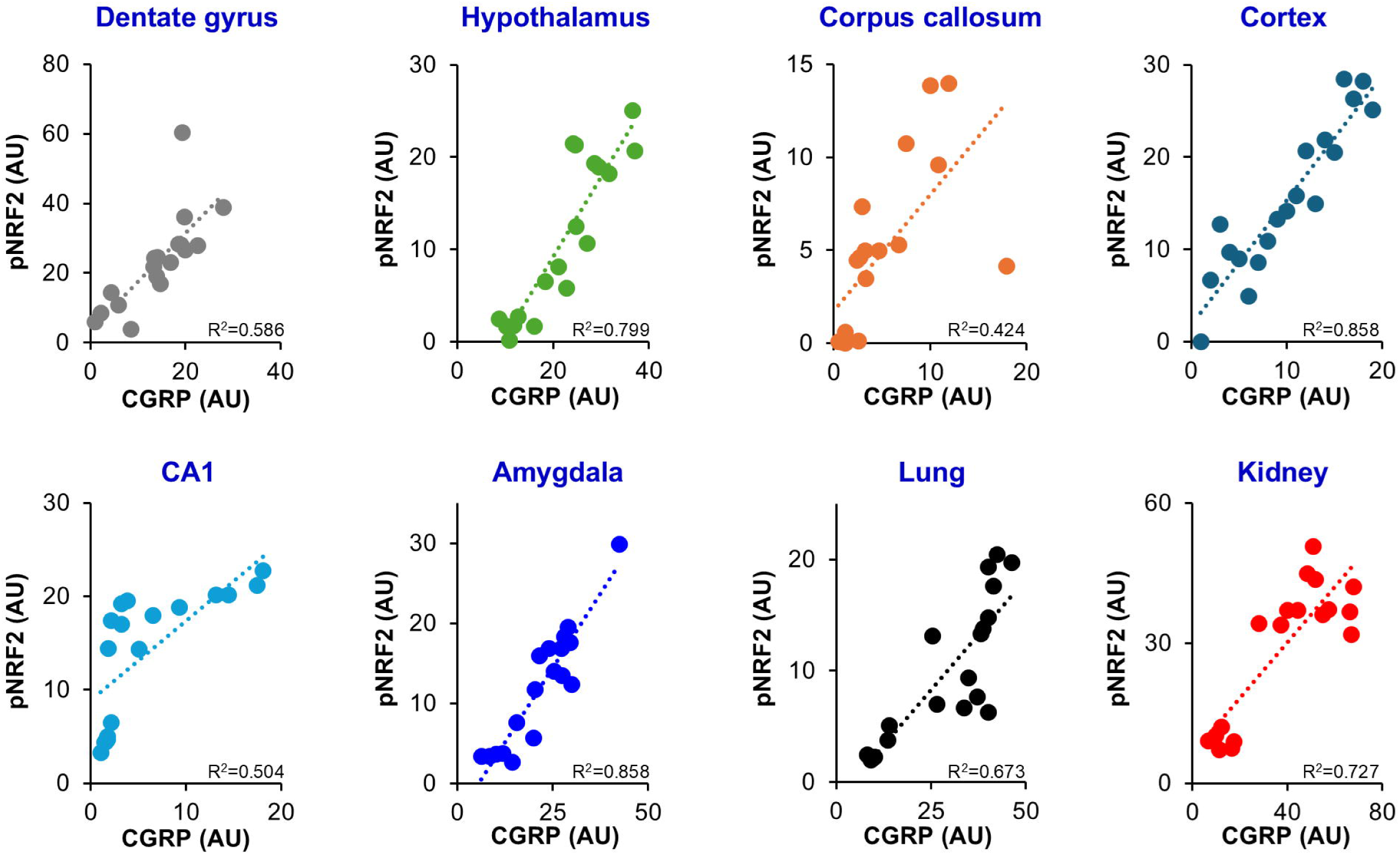

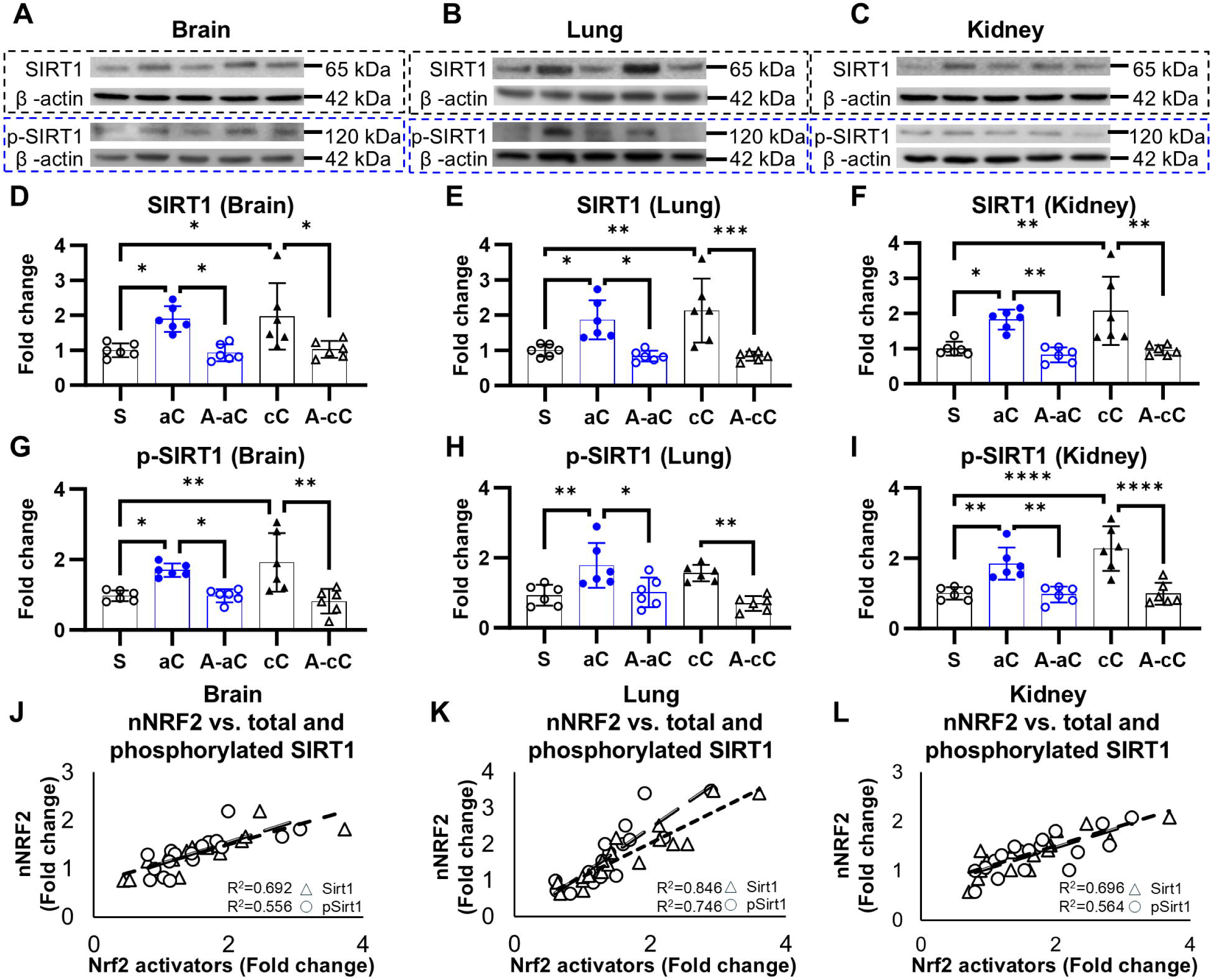

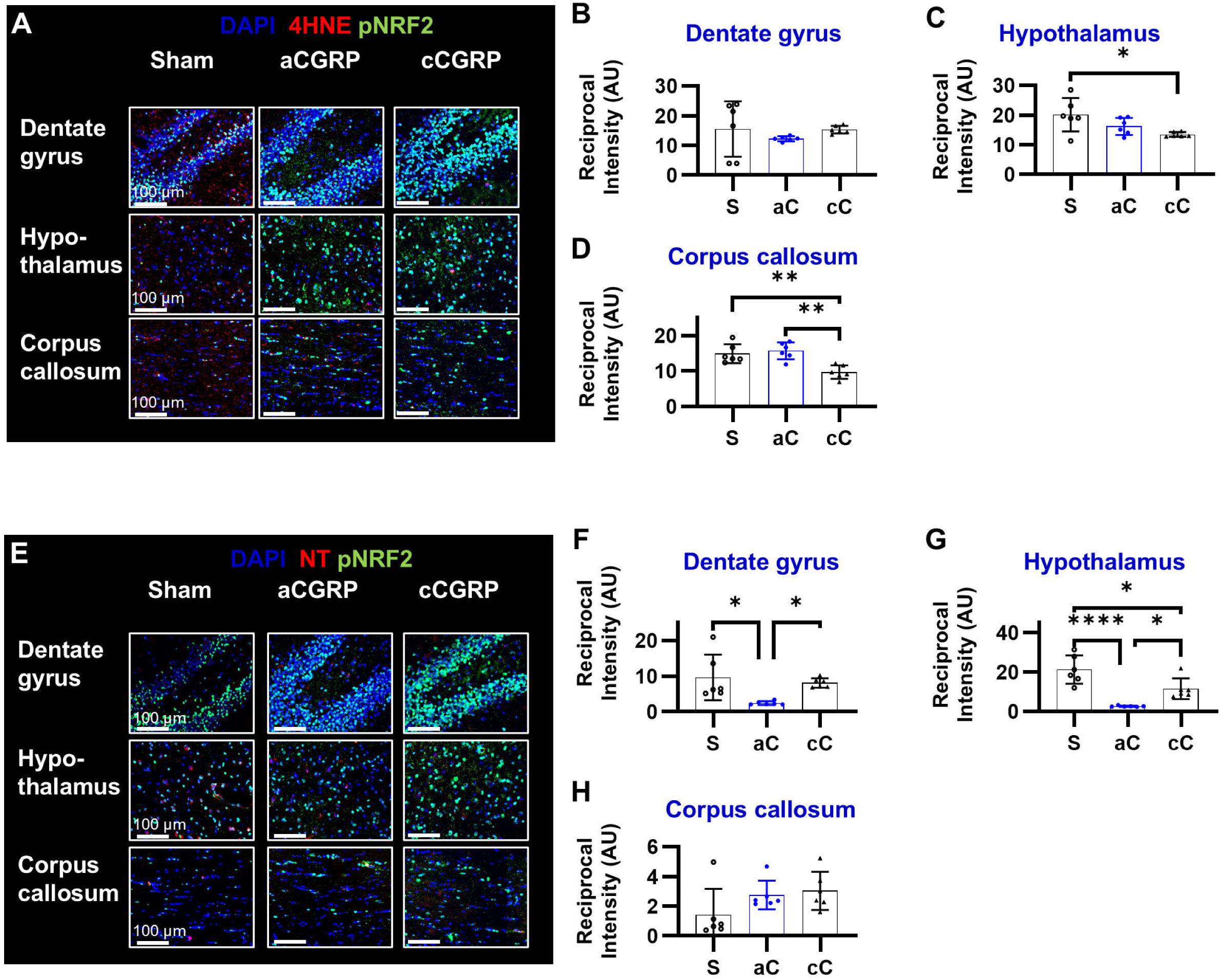

