## Supplementary Materials for "Endogenous CGRP activates NRF2 signaling via non-electrophilic mechanisms"

^ = Authors contributed to this manuscript equally

**This word file includes:**

- **Tables S1-S3**

**Table S1. Product information of commercial kits**

| **Product Name** | **Vendor** | **SKU** |
| --- | --- | --- |
| NE-PER™ Nuclear and Cytoplasmic Extraction Reagents | Thermo Scientific | 78835 |
| GSH/GSSG Ratio Detection Assay Kit (Fluorometric - Green) | Abcam | ab138881 |
| Lipid Peroxidation (MDA) Assay Kit (Colorimetric/Fluorometric) | Abcam | ab118970 |

**Table S2. The antibodies used in this study**

| **Antibodies** | **Source** | **SKU** | **Host** | **Application and dilution** |
| --- | --- | --- | --- | --- |
| CGRP | Santa Cruz Biotechnology | sc-57053 | ms | Western blotting, 1:100 in 5% milk in TBST |
| CGRP | Abcam | ab81887 | ms | Immunofluorescence, 1:50 in 1% BSA in TBST |
| NRF2 | Proteintech | 16396-1-AP | rb | Western blotting, 1:1000 in 5% BSA in TBST |
| Phosphorylated NRF2, s40 | Abcam | AB76026 | rb | Immunofluorescence, 1:100 in 1% BSA in TBST |
| AMPK | Proteintech | 66536-1-IG | ms | Western blotting, 1:1000 in 5% milk in TBST |
| Phosphorylated AMPK, T183/T172 | Abcam | AB133448 | rb | Western blotting, 1:750 in 5% milk in TBST |
| SIRT1 | Cell Signaling Technology | 8469 | ms | Western blotting, 1:1000 in 5% milk in TBST |
| Phosphorylated SIRT1, ser47 | Thermo-Fisher | 630-660 | rb | Western blotting, 1:750 in 5% milk in TBST |
| p62 | Abcam | Ab109012 | rb | Western blotting, 1:1000 in 5% BSA in TBST |
| Phosphorylated p62, ser349 | Proteintech | 29503-1-AP | rb | Western blotting, 1:1000 in 5% milk in TBST |
| KEAP1 | Proteintech | 10503-2-AP | rb | Western blotting, 1:1000 in 5% BSA in TBST |
| HO1 | Proteintech | 10701-1-AP | rb | Western blotting, 1:1000 in 5% milk in TBST |
| SOD1 | Proteintech | 10269-1-AP | rb | Western blotting, 1:2500 in 5% milk in TBST |
| NQO1 | Proteintech | 67240-1-IG | rb | Western blotting, 1:5000 in 5% milk in TBST |
| Nitrotyrosine | Santa Cruz Biotechnology | Sc-101358 | ms | Immunofluorescence, 1:100 in 1% BSA in TBST |
| 4-HNE | Abcam | ab48506 | N/A | Immunofluorescence, 1:25 in 1% BSA in TBST |
| β-actin | Sigma | A5441-.2ML | ms | Western blotting, 1:25000 in 5% milk in TBST |
| Lamin B1 | Proteintech | 66095-1-IG | ms | Western blotting, 1:5000 in 5% milk in TBST |
| DAPI Solution | Thermo Fisher Scientific | 62248 | N/A | Immunofluorescence, 1:2000 |

**Table S3. The sequences of RT-qPCR primers used in this study**

| **Genes of interest** | **Primer sequences (5’-3’)** |
| --- | --- |
| *HO1* | F:ACAGGGTGACAGAAGAGGCTAA |
|  | R:CTGTGAGGGACTCTGGTCTTTG |
| *SOD* | F:GCTCTAATCACGACCCACT |
|  | R:CATTCTCCCAGTTGATTACATTC |
| *NQO1* | F:GCGTCTGGAGACTGTCTGGG |
|  | R:CGGCTGGAATGGACTTGC |
|  | R:GGCAGGAATGGTCTCTCTCTGTG |
| *GAPDH* | F:AGGTTGTCTCCTGTGACTTC |
|  | R:CTGTTGCTGTAGCCATATTC |
