## Supplementary material for "Endogenous CGRP activates NRF2 signaling via non-electrophilic mechanisms": Whole Blot Images

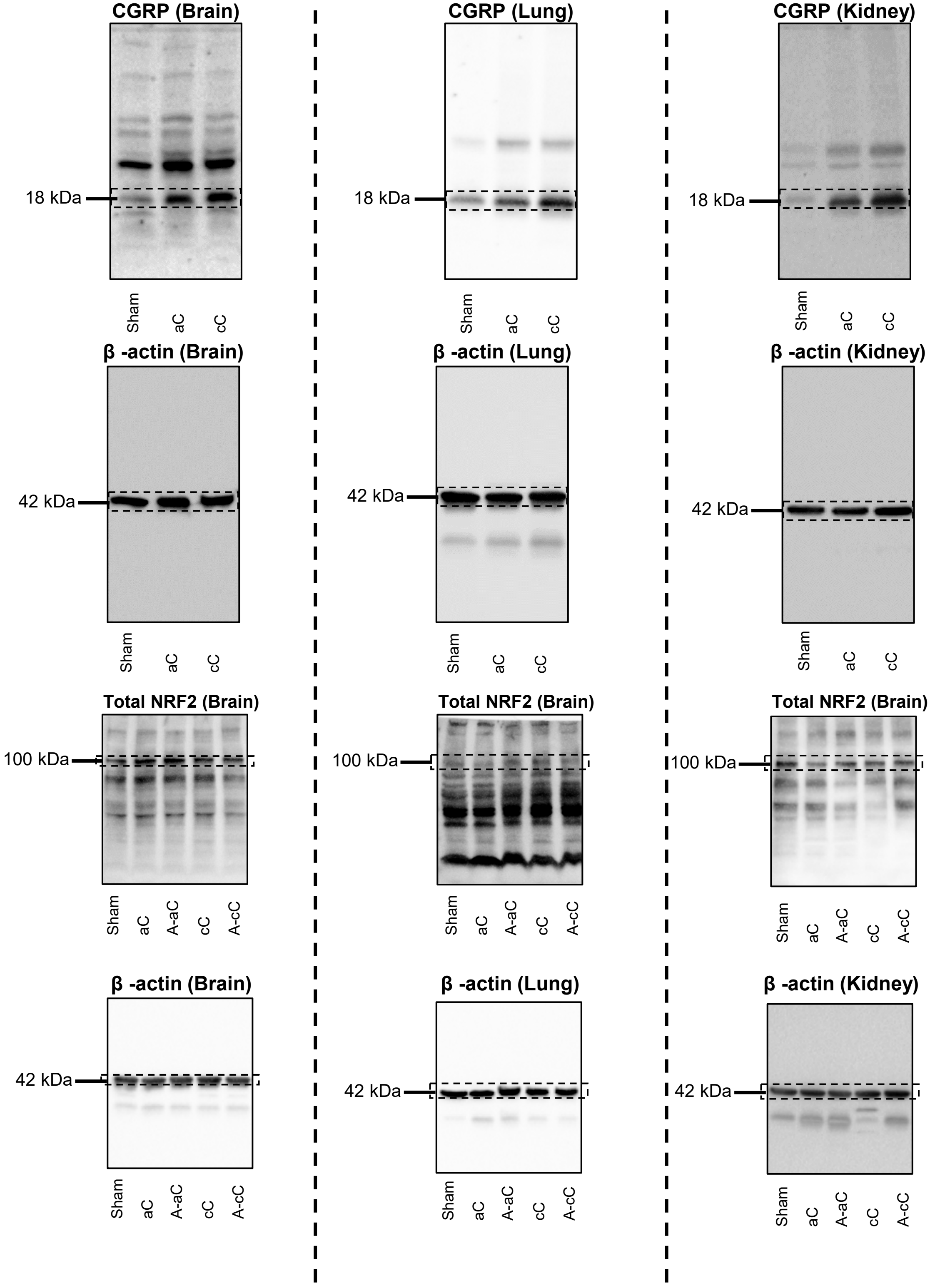


### Figure 1 – Whole Blot Images

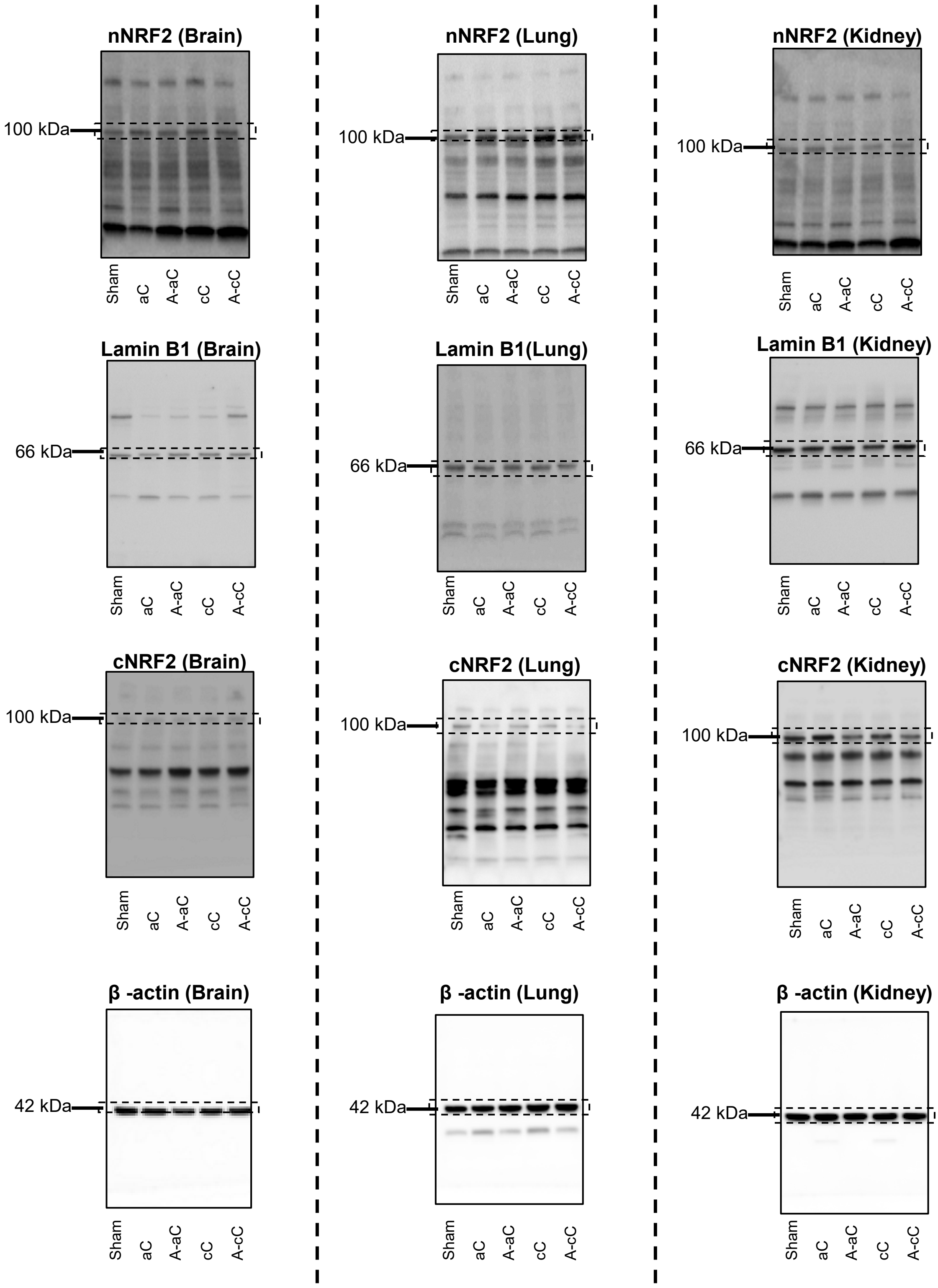


### Figure 1 – Whole Blot Images

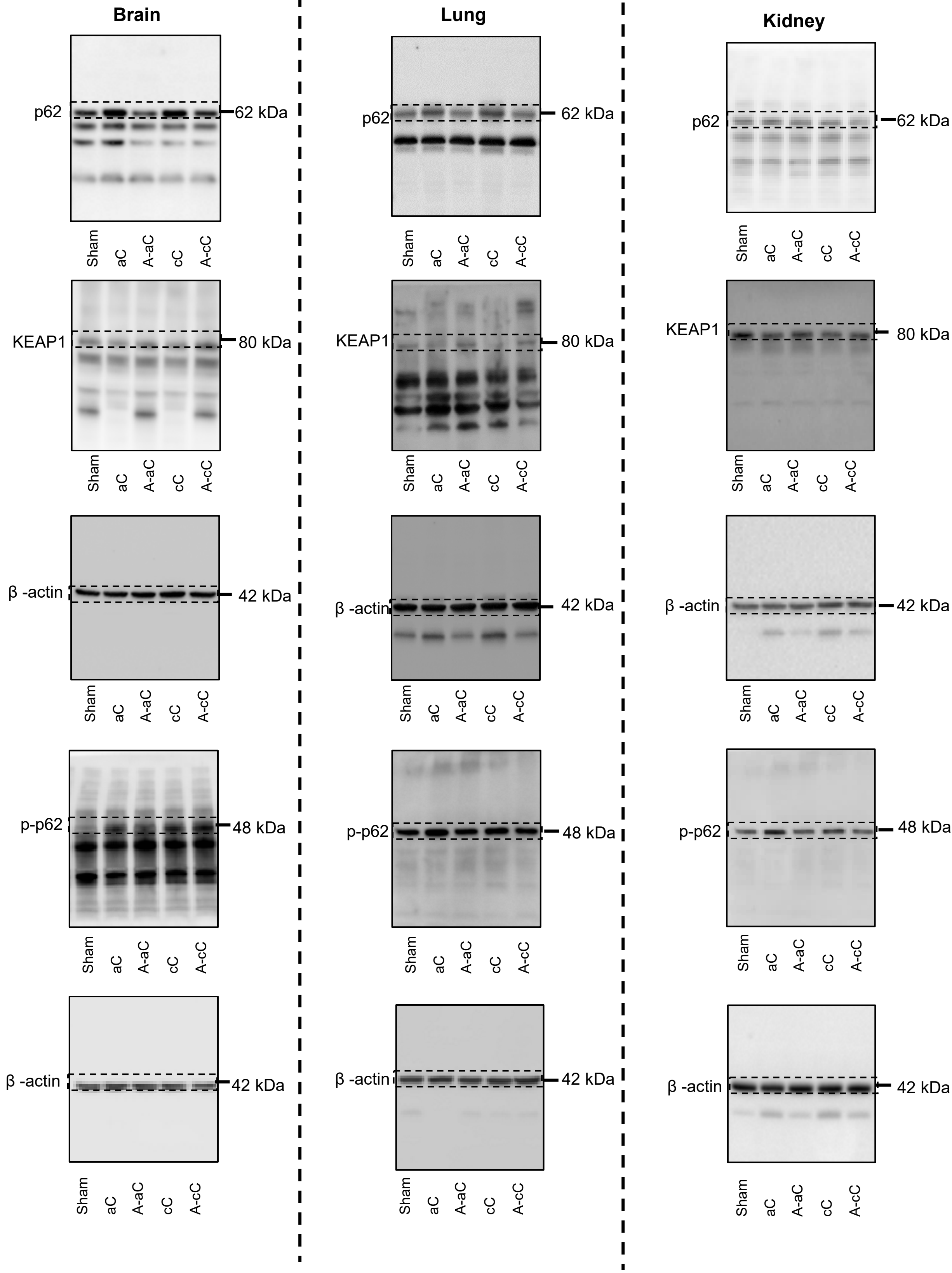


### Figure 3 - Whole Blot Images

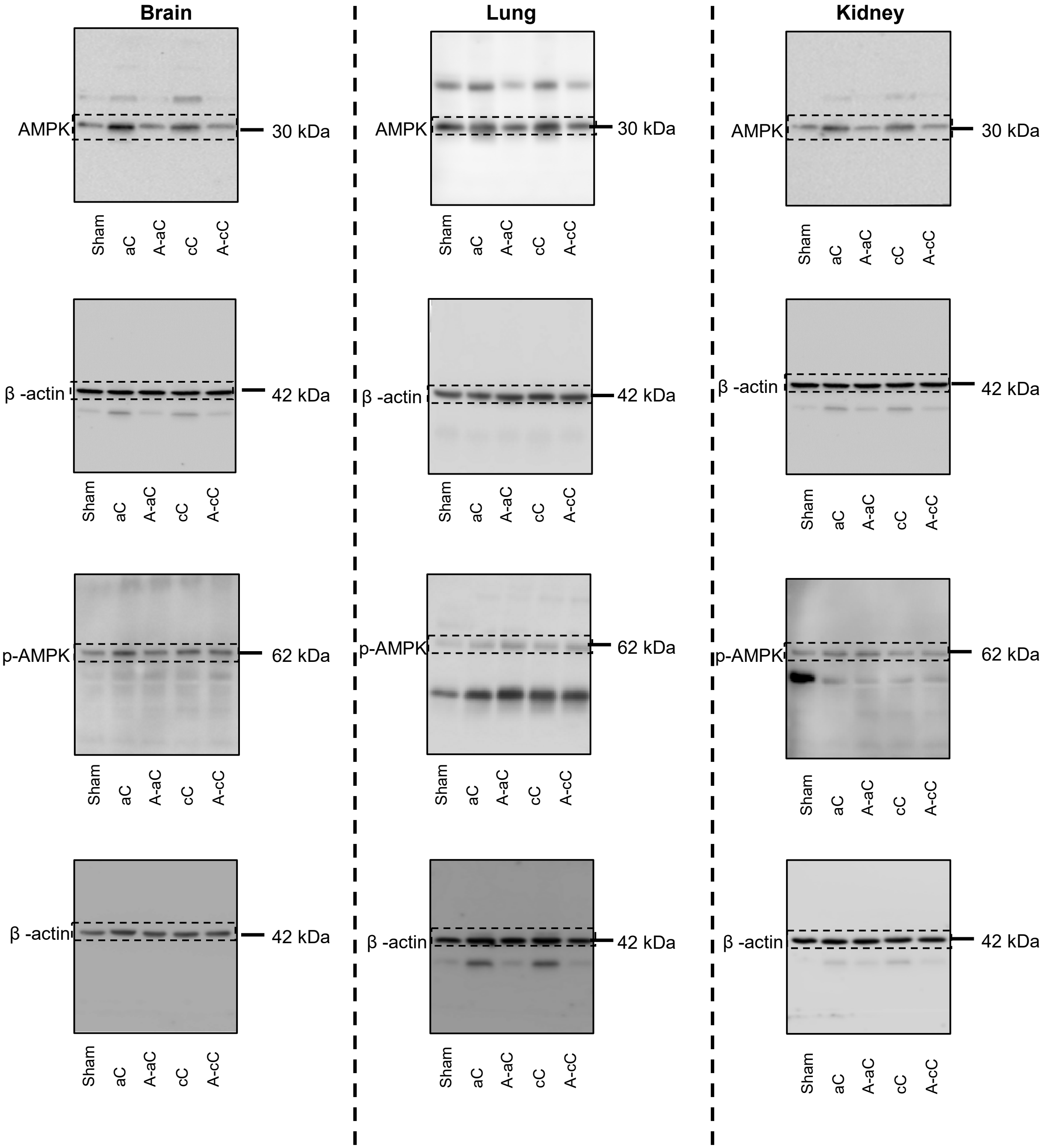


### Figure 4 - Whole Blot Images (Select β–actin shared with Supplemental Figure 6)

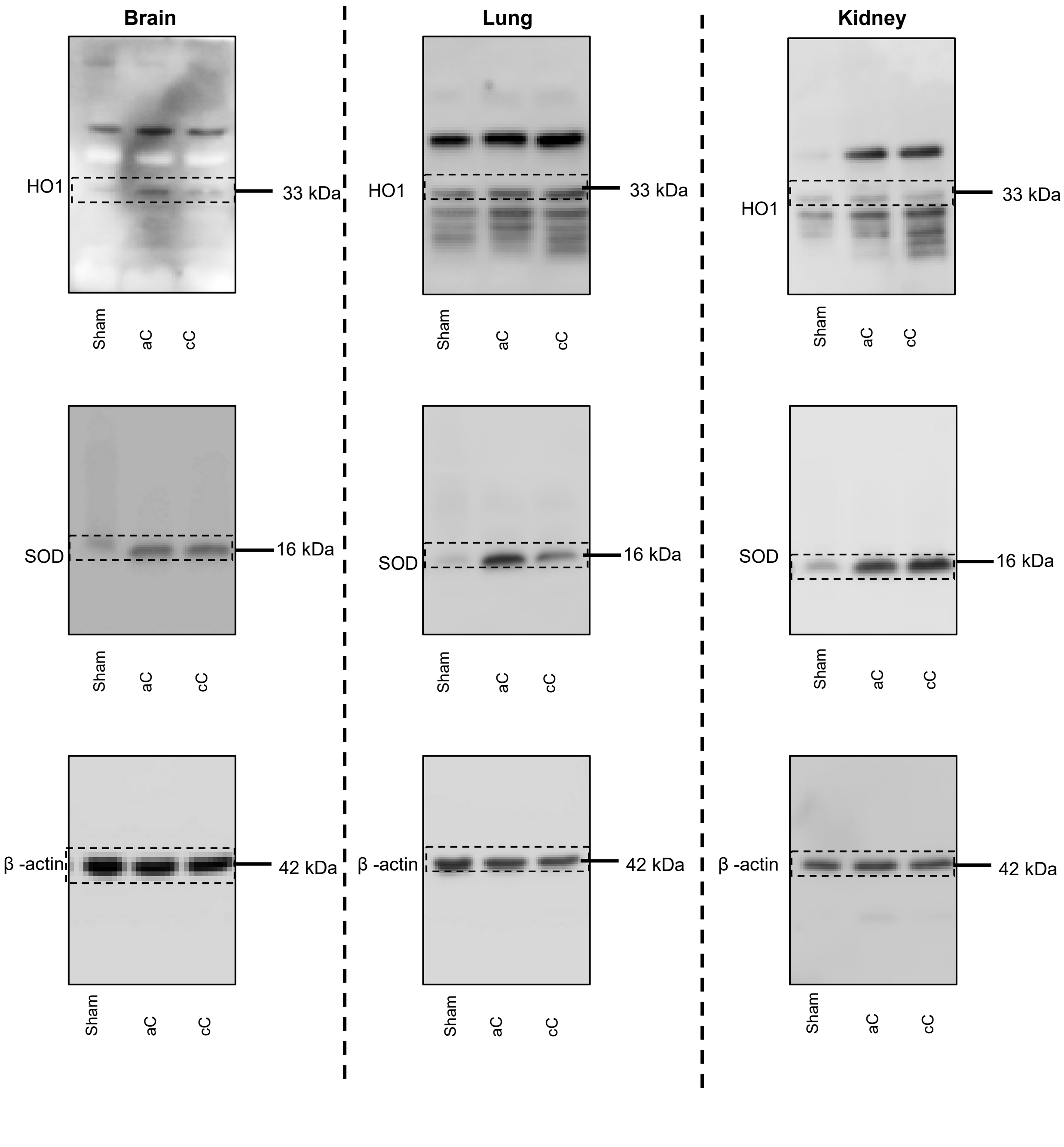


### Figure 8 - Whole Blot Images

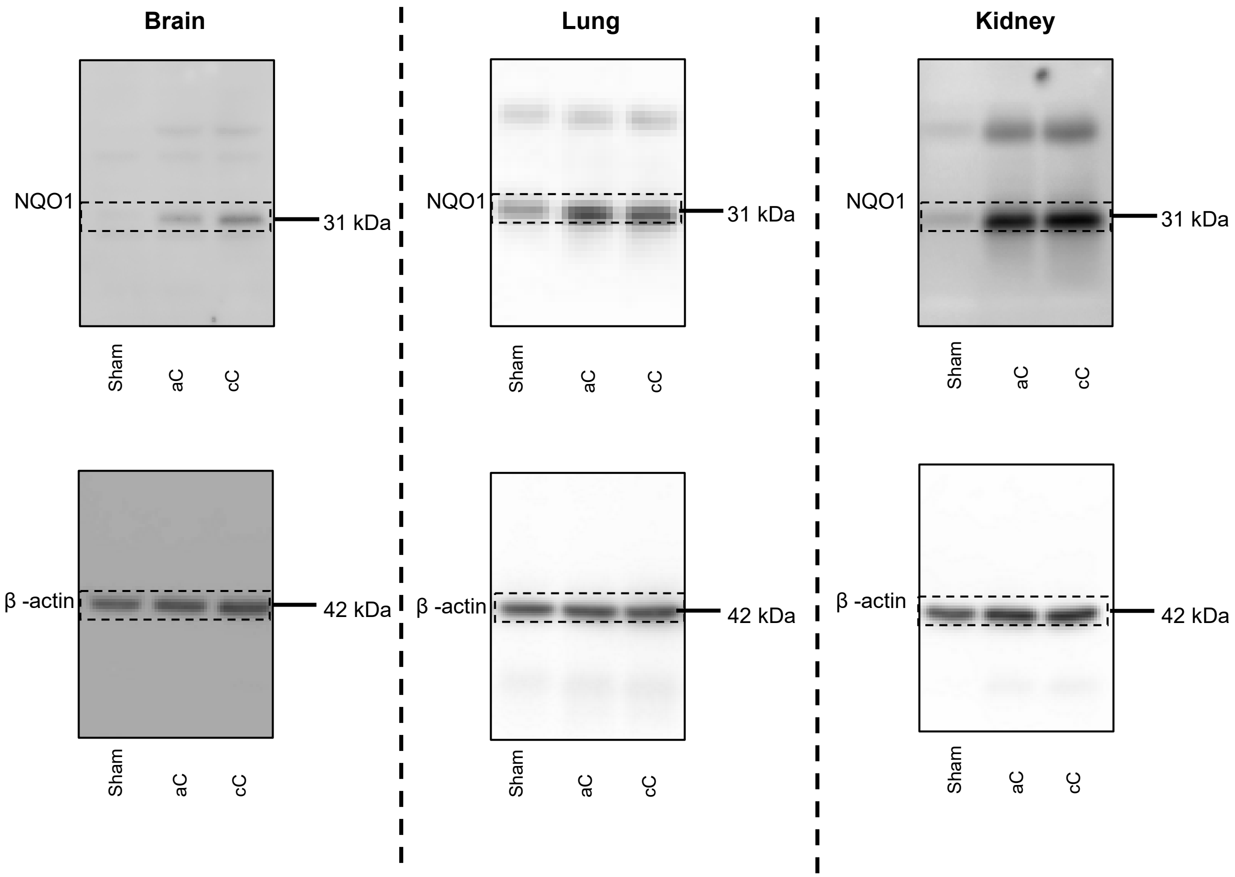


### Figure 8 - Whole Blot Images

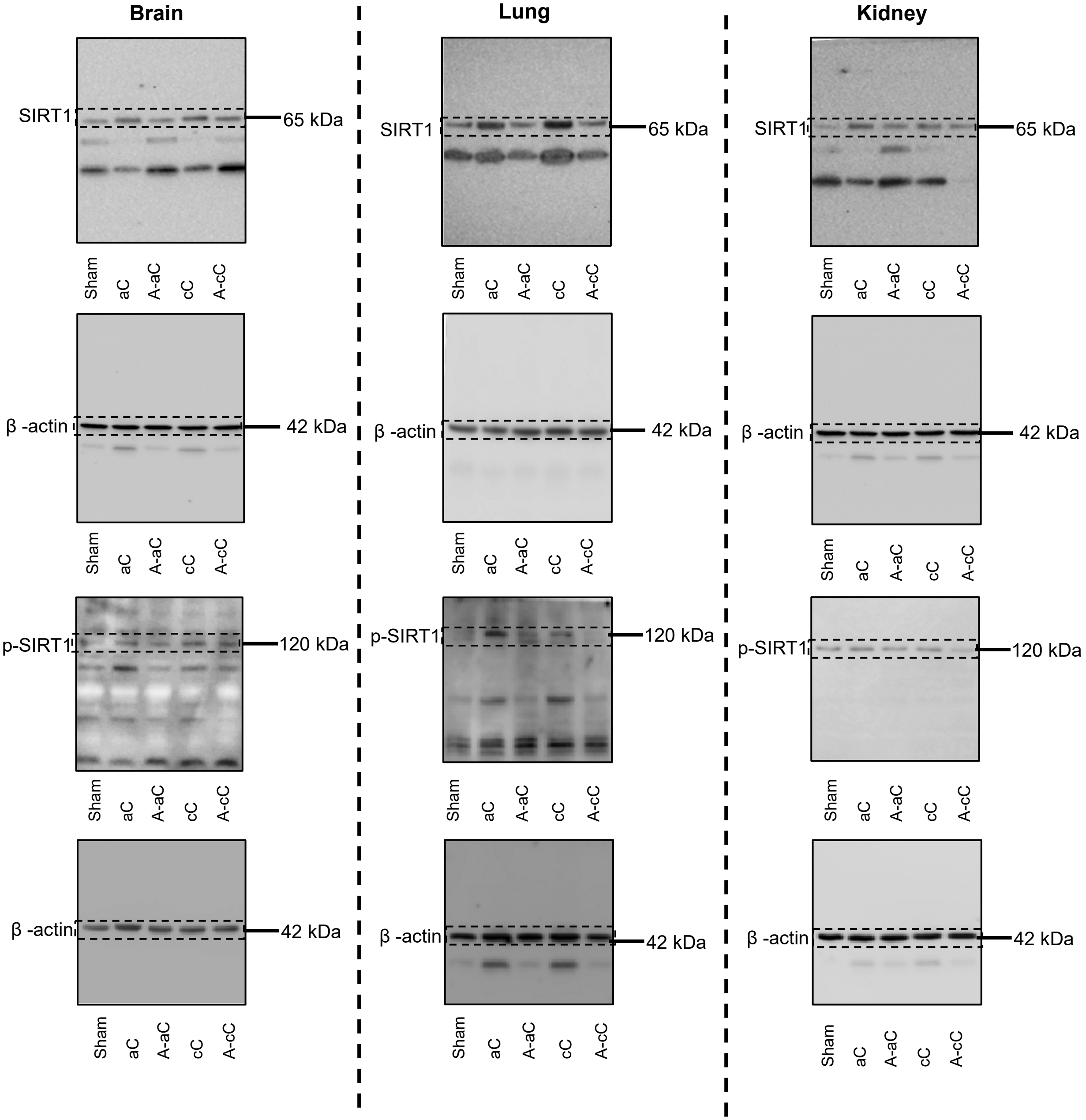


### Supplemental Figure 6
### (Select β–actin shared with Main Figure 4)
